# A cell-of-origin epigenetic tracer reveals clinically distinct subtypes of high grade serous ovarian cancer

**DOI:** 10.1101/484063

**Authors:** Pietro Lo Riso, Carlo Emanuele Villa, Gilles Gasparoni, Raffaele Luongo, Anna Manfredi, Andrea Vingiani, Annemarie Jungmann, Annalisa Garbi, Michela Lupia, Pasquale Laise, Vivek Das, Giancarlo Pruneri, Giuseppe Viale, Nicoletta Colombo, Ugo Cavallaro, Davide Cacchiarelli, Jörn Walter, Giuseppe Testa

## Abstract

High grade serous ovarian cancer (HGSOC) is a major unmet need in oncology. The persistent uncertainty on its originating tissue has contributed to hamper the discovery of oncogenic pathways and effective therapies. Here we define the DNA methylation print that distinguishes the human fimbrial (FI) and ovarian surface epithelia (OSE) and develop a robust epigenetic cell-of-origin tracer that stratifies HGSOC in FI-and OSE-originated tumors across all available cohorts. We translate this origin-based stratification into a clinically actionable transcriptomic signature, demonstrating its prognostic impact on patients’ survival and identifying novel network level dysregulations specific for the two disease subtypes.

## Introduction

Ovarian Cancer is 8^th^ most common cause of cancer death in women worldwide, being the 18^th^ most common cancer worldwide and accounting for more than 200000 new cases every year^1^. Its most prevalent form is high grade serous ovarian cancer (HGSOC) which accounts for more than 70% of cases and is usually diagnosed at already advanced stages being mostly asymptomatic earlier on^2^. In the 30 years since the introduction of carboplatin-based regimens, the cure rate has changed only negligibly, largely due to the precocious dissemination (favored by the anatomic continuity with the abdominal cavity) and the dearth of physiopathologically meaningful models that can recapitulate the pathogenesis and progression of the human disease, an increasingly recognized need for the field as a prerequisite for the identification and validation of new therapeutic targets^3,4^. In turn, the lack of actionable human models is due to a significant extent to the persistent uncertainty regarding the cell of origin of HGSOC, with the two candidate originating tissues identified in the distal tract of the fallopian tube (fimbrial epithelium, FI) and the epithelial lining of the ovary (ovarian surface epithelium, OSE)^5^. In recent years, growing evidence has pointed to the FI as the main source of HGSOC, starting from prophylactic salpingo-oophorectomy specimens from patients with increased risk of ovarian cancer, which revealed how the fimbria is frequently hit by pre-cancerous lesions referred to as serous tubal intraepithelial neoplasia (STIC). This suggested that lesions labelled as ovarian tumors could actually result from seeding of primary fimbrial tumors, through trapping of STIC-derived cells in the lumen of the ovary as inclusion cysts, favored by the ruptured stigmas of the surface epithelium during the menstrual cycle. Initial transcriptomic and methylomic analyses also uncovered higher similarity between HGSOC and FI with respect to the OSE, supporting a non-primarily ovarian origin^6,7^. Mutational analysis of STIC, primary HGSOC and peritoneal metastasis from the same patient revealed the presence of a shared mutational spectrum, thus reinforcing the notion of a primary of not exclusive fallopian origin^8^. These lines of evidence notwithstanding, other convergent sources failed to settle this fundamental developmental question, starting from the observation that in a significant proportion of HGSOC samples no STIC precursor lesion can be identified. This indicates that the fimbrial origin seeding model cannot be universally applied to all HGSOC samples. Indeed, the reverse route of dissemination has been also shown to be equally plausible, with genomic studies supporting the possibility that at least a fraction of STIC actually represent metastatic lesions of primarily ovarian lesions, a model underscored by the experimental finding that HGSOC-derived spheroids can implant into the fallopian tube epithelium^9^. While it is yet to be clarified whether also early OSE lesions can seed onto the fimbrial surface and give rise to STIC, there thus remains fundamental uncertainty about the developmental origin of HGSOC, both in terms of the general distribution between FI or OSE origins and, more importantly, in terms of patient-tailored assays to assign developmental origin on a case-to-case basis.

Here we present a novel approach to solve this problem, grounded in the tumor retention of a cell of origin-specific DNA methylation print (OriPrint) that allows to stratify HGSOC in FI-and OSE-originated tumors (FI-like and OSE-like) robustly across all available datasets. We show that both epithelia serve as *bona fide* origin for this disease and that the cell of origin explains most of the variance existing among tumors. We then translate these findings into a transcriptional readout through a particularly cost-effective, clinically-relevant transcriptomic analysis. This revealed the prognostic value of a cell of origin-based classifier in an independent, well-characterized retrospective cohort. Specifically, we found that OSE-like tumors carry a significantly worse prognosis that can be ascribed to a reduced inflammatory response, coupled to increased survival and cell-to-cell signaling in this specific tumor subset. Finally, our deconvolution of developmental origin uncovers new genes and pathways specific for OSE-like HGSOC that open new strategies for an improved management of HGSOC patients.

## Results

DNA methylation is one of the most characterized epigenetic mark, playing a critical role in chromosome replication and regulation of gene expression. Its tightly regulated propagation and deposition at specific loci, ensures both the inheritance over multiple cell divisions of gene regulatory features and the activation/repression of gene regulatory programs in response to stimuli. Thus, it is not surprising that mutations/alterations occurring on key regulators of the DNA methylation machinery can lead to hypo/hypermethylation at multiple loci, promoting tumorigenesis^10^. Still, despite a frequently diffused resetting of the methylomic landscape, recent evidence suggests that a portion of DNA methylation prints can be retained during tumor evolution, a process resembling the so-called epigenetic memory which has been extensively studied in the developmental setting^11^. Indeed, analyzing the DNA methylome of tumor cells has allowed to track tumor clonal evolution, to an extent comparable to genetic fingerprinting, with converging trajectories along tumor development^12^. Importantly, the retention in tumor cells of epigenetic prints of the tissue of origin has also allowed to associate the tissue of origin to cancers of unknown primary (CUP)^13^, allowing for the first time a tailored treatment and improved care for patients. Also, the definition of tumor-specific DNA methylation traits allowed to re-classify central nervous system tumors, impacting routine diagnostics for these diseases^14^.

In order to probe whether DNA methylation could be used as a developmental tracker for HGSOC’s cell of origin, we derived genome-wide DNA methylation profiles of short-term cultures of fimbrial epithelium (FI, n = 11), ovarian surface epithelium (OSE, n = 7), solid tumor-derived (HGSOC, n = 11) and ascites-derived (AS, n = 13) tumor cells. We checked in our dataset as well as in two independent published datasets generated from frozen tissues^7,15^(Omaha and Melbourne cohorts) whether the global variance in DNA methylation could already stratify tumors on the basis of their cell of origin. To this end we used Uniform Manifold Approximation and Projection (UMAP) visualization, a non-linear dimensionality reduction technique that preserves the global structure of the data^16^, to capture the largest fraction of variability in DNA methylation. We found that neither in published datasets nor in our cohort it was possible to bipartition tumor samples according to FI or OSE global methylation (Supplementary Figure 1), since most of the variability is driven by the differences between normal and tumor samples. We thus reasoned that given such a distribution of variance, we could enhance the potential sensitivity of DNA methylation as a cell-of-origin tracker by first defining the specific subset of differentially methylated CpG able to distinguish FI and OSE. We thus identified 13926 differentially methylated sites (DMS) between the two normal cell types in culture (Figure 1A) (henceforth OriPrint), which mapped preferentially in open sea^17^ CpG areas and were functionally annotated mostly to introns and intergenic regions (Supplementary Figure 2). Next, we determined whether this blueprint, which was derived from cells in culture, was able to segregate correctly FI and OSE tissues (Figure 1B). As shown in Figure 1C, a hierarchical clustering based on Pearson’s correlation confirmed that the tissue samples were correctly divided into FI and OSE, thus proving that our culture conditions recapitulated the complexity and preserved the essential epigenetic properties of tissues *in vivo*. Indeed, the very same FI sample form the Omaha cohort that already been misclassified as OSE in the original published dataset^7^ behaved in the same manner also in our classification (Figure 1C), underscoring the robustness and sensitivity of our classifier.

**Figure 1.**
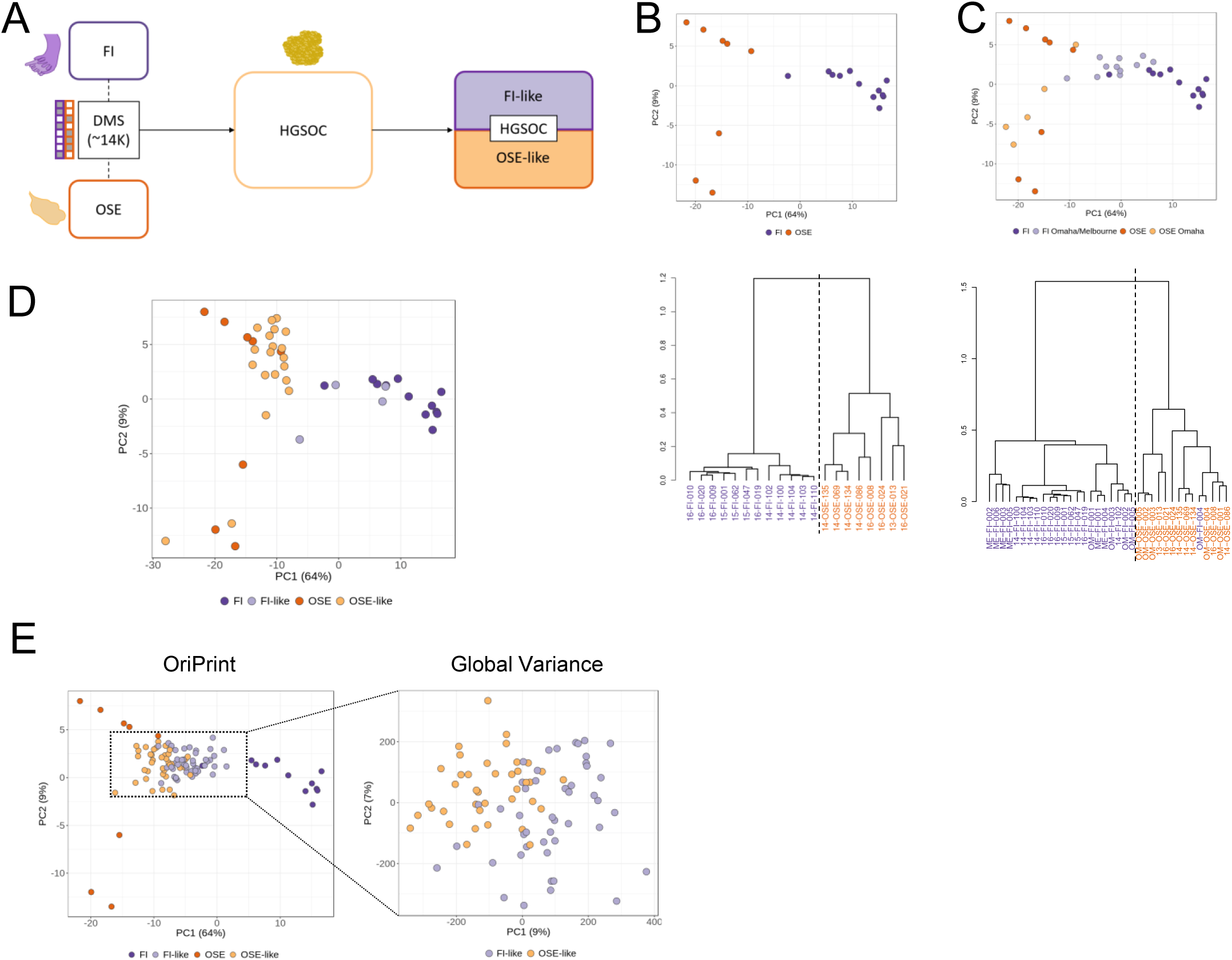
OriPrint is able to stratify HGSOC on the basis of its cell of origin. **A.** Schematic representation of the experimental pipeline. FI = Fimbrial Epithelium; OSE = Ovarian Surface Epithelium; HGSOC = High Grade Serous Ovarian Cancer; FI-like = tumors originating from Fimbrial Epithelium; OSE-like = tumors originating from Ovarian Surface Epithelium. **B.** *Top:* PCA analysis of FI and OSE samples from IEO cohort (purple and orange, respectively) considering ORIPrint CpGs; *Bottom:* Hierarchical clustering of the same samples, distance = Pearson’s Correlation. **C.** *Top:* PCA analysis of FI and OSE samples (tones of purple and orange, respectively) from IEO, Omaha and Melbourne Cohorts considering OriPrint CpGs in the space defined by normal samples. *Bottom:* Hierarchical clustering of the same samples, distance = Pearson’s Correlation. **D.** PCA analysis of normal and tumor samples from IEO cohort, annotated using Pearson’s correlation-based classification in the space defined by normal samples. **E.** PCA analysis of normal samples from IEO cohort and tumor samples from Melbourne Cohort, annotated using Pearson’s correlation-based classification both in the space defined by OriPrint and normal samples (left) and by the whole set of CpGs (right).

**Figure 2.**
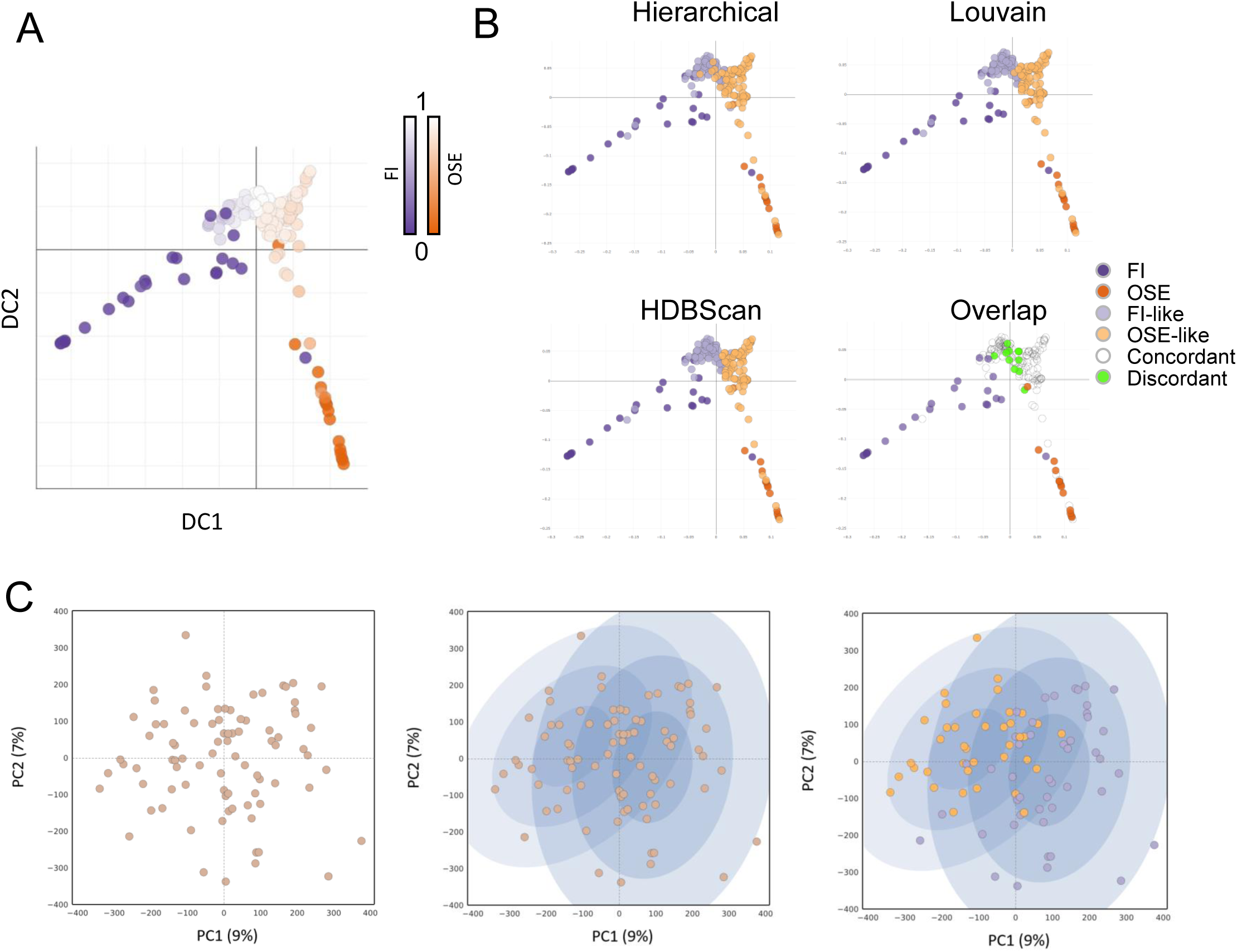
OriPrint is a solid stratifier and establishes the tissue of origin as a major source of variance for HGSOC. **A.** Diffusion map with pseudotime timeline performed on OriPrint CpGs for samples of all cohorts. The origin is situated either in the distal FI (purple origin) or OSE (orange origin) samples. **B.** Diffusion maps showing the classification output for the three indicated clustering methods. The overlap plot shows in white the samples that are concordantly classified by all three methods and in green the samples that have a different classification in at least one of the three methods. **C.** PCA analysis coupled to Gaussian Mixture Model (GMM) clustering of Melbourne tumor cohort. Left: first two components of global variance in DNA methylation for the considered samples; Middle: the two probability distributions calculated by GMM; Right: Overlay of the OriPrint classification, showing a consistent overlap with G’s distributions.

To see whether OriPrint could be used to stratify tumor samples, we applied the same methodology to our cohort of tumor samples and showed that tumors were bipartitioned through the OriPrint in two classes that we define as FI-like and OSE-like tumors (Figure 1D, Supplementary Figure 3).

**Figure 3.**
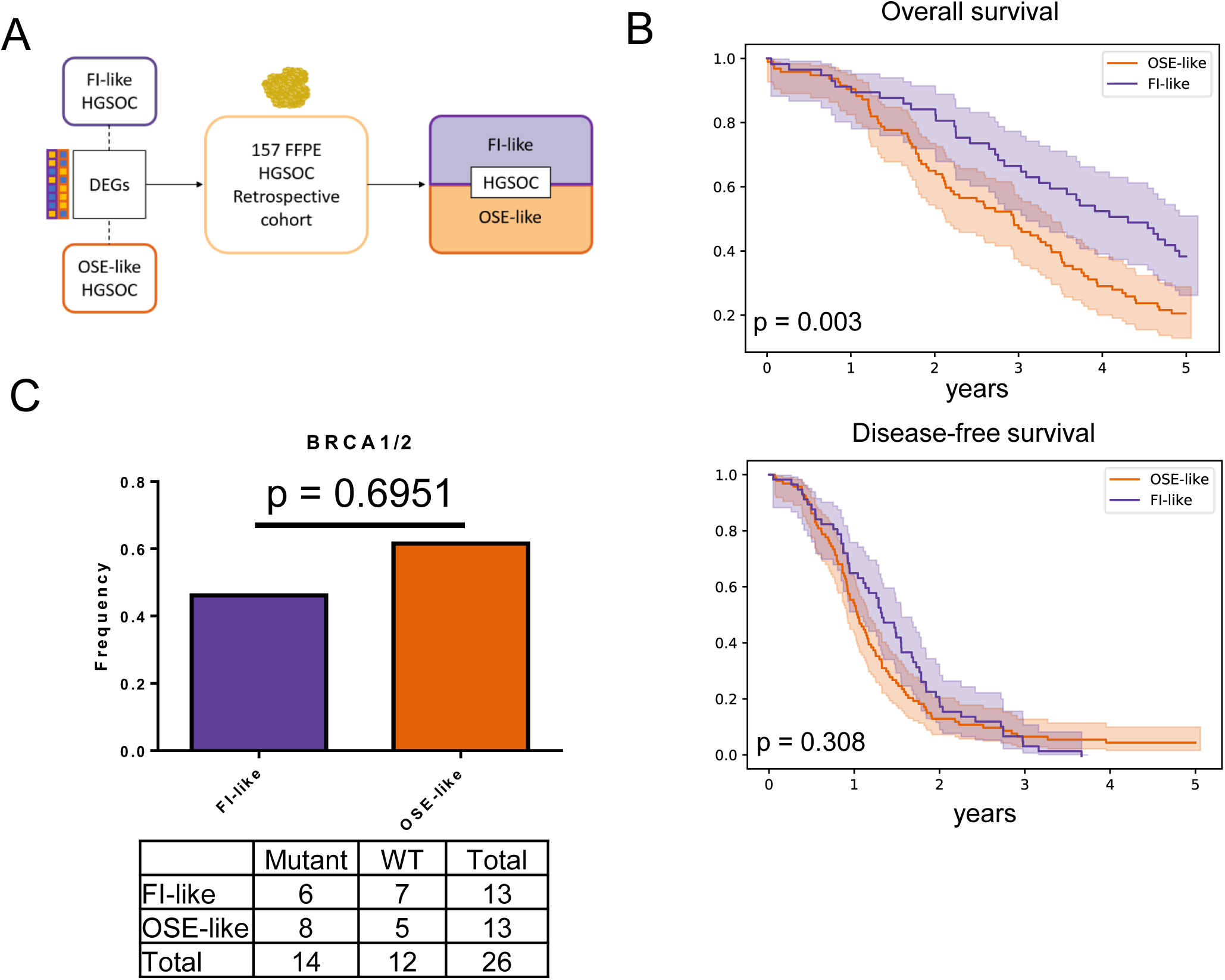
The cell of origin of HGSOC impacts the prognosis of patients independently of BRCA1/2 mutations. **A.** Schematic representation of the strategy based on RNAseq to stratify a retrospective cohort of HGSOC. **B.** Overall (top) and disease-free (bottom) survival of patients stratified by the cell of origin of HGSOC. Light-colored areas represent confidence intervals. **C.** Mutational Status of BRCA1/2 in the retrospective cohort classified in FI-like (purple) and OSE-like (orange) tumors, shown as mutational frequency (top barplot) and contingency table (bottom). isher’s Exact Test was performed based on the contingency table.

Next, to verify that our approach could be translated to independent tumor cohorts as well as to frozen biopsies, we applied the OriPrint classifier to the Melbourne cohort and analyzed the global variance in DNA methylation across FI-like and OSE-like HGSOC. Interestingly, by principal component analysis we could show that the superimposition of the categories defined by OriPrint identified two clear clusters in the first two components of variance, thus indicating that the cell of origin is the first responsible of the variability existing in DNA methylation between HGSOC samples (Figure 1E).

Together these results show that OriPrint is able to robustly stratify HGSOC tumors into FI-like and OSE-like subtypes across independent clinical cohorts and sample processing pipelines. To further confirm this finding, we used diffusion map coupled to pseudotime analysis, that was previously used for single cell RNA sequencing data to derive the developmental progression of cells and identify branching decisions and differentiation endpoints^18,19^, to highlight whether an evolutionary timeline exists between tumors and the two origins. Using OriPrint and setting the origin in both OSE and FI samples, we calculated the two pseudotime lines and could show that HGSOC samples are a mandatory step in pseudotime evolution between the two normal tissues. Moreover, we showed that the intersection between the two paths occurs centrally in the tumors’ distribution (Figure 2A), thus further confirming that both origins are plausible for HGSOC. Instead, using the whole set of CpGs, we could not score any clear evolution between FI and OSE and HGSOC (Supplementary Figure 4), thus proving that OriPrint is necessary to stratify tumors according to their cell of origin.

**Figure 4.**
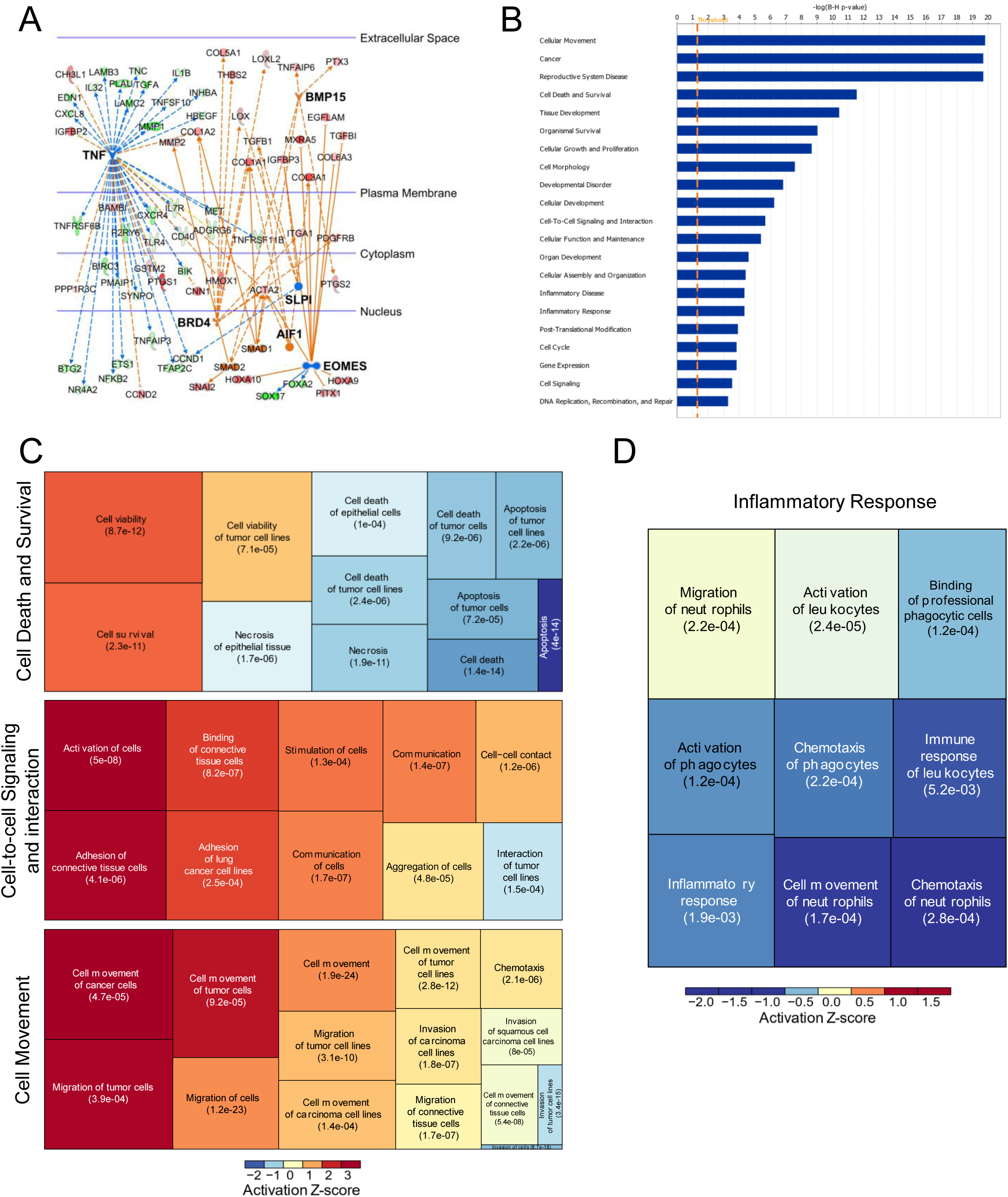
Gene expression pattern of OSE-like tumors reveal a lower inflammatory response coupled to increased survivability and active cell-to-cell signaling. **A.** IPA causal network analysis performed on WGCNA eigengenes associated with OSE-like tumors. Blue: regulator genes whose pathway is predicted to be inhibited; Orange: regulator genes whose pathway is predicted to be activated; Red: Upregulated eigengenes; Green: Downregulated eigengenes **B.** IPA Disease and Function Enrichment Analysis on WGCNA eigengenes associated with OSE-like tumors. Enrichment p-values are shown after Benjamini Hochberg FDR correction. **C.** Treemap of three of the categories in (B). Boxes dimension is derived on activation Z-score. Enrichment p-values are shown as in B. **D.** The Inflammatory Response IPA Disease and Function Enrichment Analysis Category predicted to be inhibited in OSE-like vs FI-like tumors.

In order to determine whether the classification was solid and not bound to the clustering method used, we applied two additional clustering methods, Louvain^20^ and HDBScan^21^, to stratify HGSOC and checked the concordance of classification by OriPrint. We could show that, overall, we could achieve > 92% concordance of classification among the three clustering methods used, thus proving the robustness of this approach to define FI and OSE-originated tumors (Figure 2B).

Next, we sought to determine whether by this method the same conclusions could be drawn directly from the full set of CpG or if OriPrint was necessary to define the classification into FI-like and OSE-like tumors. We applied Gaussian Mixture Model (GMM) clustering to the entire set of CpG in the space derived from the first two components of variance (i.e. the components accounting for the highest variance in the system) and were able to derive two probability distributions for samples. We then overlaid the classification derived from OriPrint and could show that the two clustering methods are mostly superimposable (Figure 2C). Thus, we could show that HGSOC cell of origin is one of the main determinants of the differences in DNA methylation existing among HGSOC samples.

### Stratification of a retrospective HGSOC cohort

In order to check whether the two subtypes had a different impact on patients’ prognosis we decided to stratify a well characterized FFPE cohort of HGSOC (n = 151 independent patients) (Supplementary Table 1). Since FFPE fixation compromises bisulfite-based DNA methylation analysis we concentrated our effort on analyzing transcriptomic profiles that we obtained from our cohort of samples in culture and from macrodissected FFPE tissues. We then used the differentially expressed genes between FI-like and OSE-like tumors that we previously identified through DNA methylation (n=3 FI-like, n=8 OSE-like) to stratify the cohort and to perform survival analysis (Figure 3A). Using this method, we found that the patient’s affected by OSE-like tumors had a worse prognosis, as assessed by overall survival analysis (difference in survival 1.7 years, p value < 0.005). Instead, we could not score an impact on the disease-free survival of patients (Figure 3B), indicating that both subtypes are equally eager to recurrence *in vivo.*

**Table 1.**
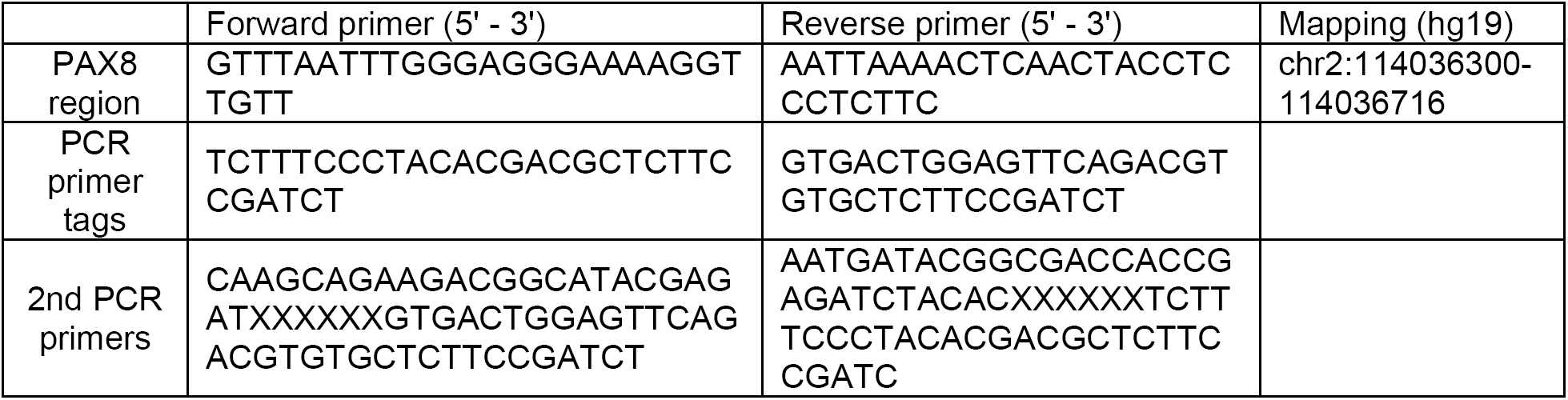
Primer sequences used in this study. ‘X‘ refers to sample-specific barcode sequences.

To check whether the difference in overall survival was due to the presence of subtype-specific mutations, we analyzed the mutational status of BRCA1/2 in our samples, that has been shown to positively correlate with patient’s survival^22^. We could show that BRCA1/2 mutations are present in both subtypes with no difference in frequency in the two groups, thus indicating that the mutational status of these genes is not responsible for the difference in survival existing between the two groups (Figure 3C).

### Definition of subtype specific transcriptional signatures

To gain insight in the specific transcriptional features that characterize the most detrimental HGSOC, i.e. OSE-like tumors, we performed Weighted Correlation Network Analysis (WGCNA) on all sample categories (FI: 6, FI-Like: 3, OSE: 8, OSE-Like: 8 samples) and identified 1378 genes whose behavior correlated in OSE-like tumors.

In order to identify the upstream regulator pathways that could account for the specific gene expression pattern of these tumors, we performed IPA causal network analysis on WGCNA genes. Through this approach we identified 6 pathways that were regulating 80 genes specific for OSE-like tumors (TNF, BRD4, EOMES, BMP15, AIF1, SLPI). More in detail, the TNF pathway is predicted to be deactivated in OSE-like HGSOC, suggesting a reduced inflammatory response for these tumors, while the other pathways are predicted to be activated and associated to extracellular matrix remodeling and the WNT-β-catenin pathway (Figure 4A).

To understand the downstream effect of WGCNA genes on OSE-like tumors’ fitness, we performed IPA disease and function analysis. We highlighted among the other categories a consistent inactivation of the cell death related categories, with a concomitant activation of the cell survival-related genes, and the activation of categories related to cell-to-cell signaling and interaction and cell movement (Figure 4B-C). Taken together, these data suggest a higher fitness and survivability of OSE-like tumors.

In order to investigate the effect related to the alteration in the identified pathways in FI-like and OSE-like tumors, we performed differential expression analysis between these two subtypes.

We found 146 differentially expressed genes that distinguished the two categories, consistently associated with decreased inflammatory response in OSE-like tumors, thus confirming our observations in WGCNA (Figure 4D).

To further dissect the molecular features that characterize these two tumor subtypes, we took advantage of the published molecular classification for HGSOC by Tothill and colleagues^23^ that identified four molecular subclasses (C1 – mesenchymal, C2 – immunoreactive, C4 – proliferative, C5 – differentiated).

We used the minimal signature of validated classifier genes^24^ that were expressed in our dataset and evaluated whether we could assign any of the signatures to our stratified tumors.

Interestingly, OSE-like tumors molecularly resembled mesenchymal tumors, while skewing away from the immunoreactive phenotype (Figure 5A-B).

**Figure 5.**
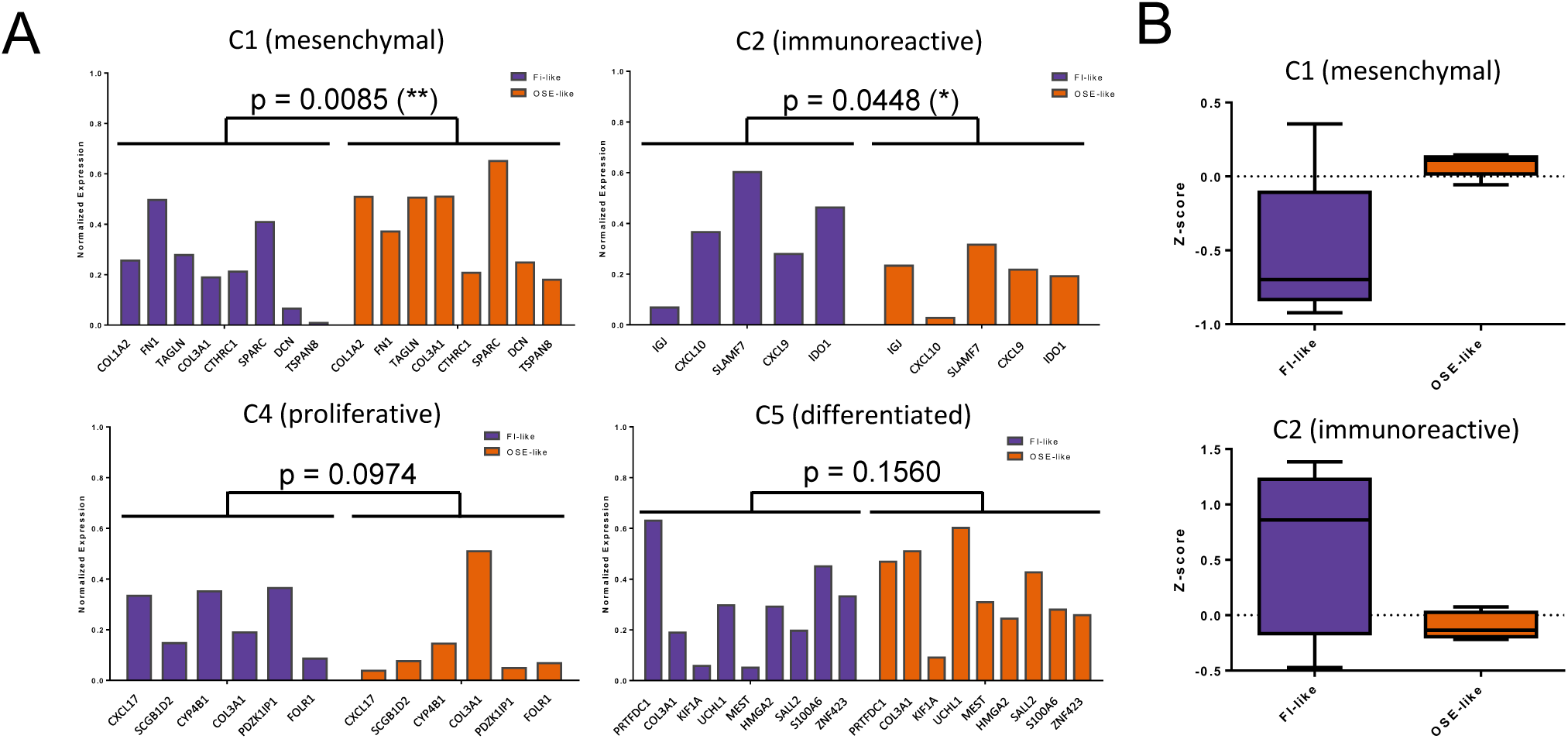
OSE-like tumors have a mesenchymal, non-immunoreactive molecular phenotype. **A** Barplot of the mean expression levels of HGSOC molecular signatures in FI-like (purple bars) and OSE-like (orange bars) tumors. Two-way ANOVA analysis was used to calculate the significance for the difference in expression of the signature in the two groups. **B.** Boxplot of the z-score relative to the expression of the mesenchymal (top) and immunoreactive (bottom) signatures in FI-like (purple) and OSE-like (orange) tumors.

### PAX8 is differentially regulated according to the origin of HGSOC

PAX8 has been historically described as a defining marker for HGSOC^25,26^, whose expression is shared with the fimbrial epithelium. This latter feature has been proposed as one of the demonstrations of the Müllerian origin of HGSOC. Since, not all tumor samples express PAX8, we sought to determine whether PAX8 expression could be related to the cell of origin of this disease.

We indeed found that PAX8 is expressed in FI but not in OSE samples and remains differentially expressed between FI-like and OSE-like tumors, following the same expression pattern of the tissues of origin. To understand whether this pattern of expression was reflected also at the regulatory level, we analyzed the level of methylation of its promoter in our samples. Consistent with the differences in gene expression, PAX8 was differentially methylated among FI and OSE samples and also between FI-like and OSE-like tumors, in an anticorrelative fashion with gene expression (Figure 6A-B). We validated these findings through qPCR and targeted bisulphite sequencing (Supplementary Figure 5), thus confirming a differential regulation for this gene according to the cell of origin of this disease.

**Figure 6.**
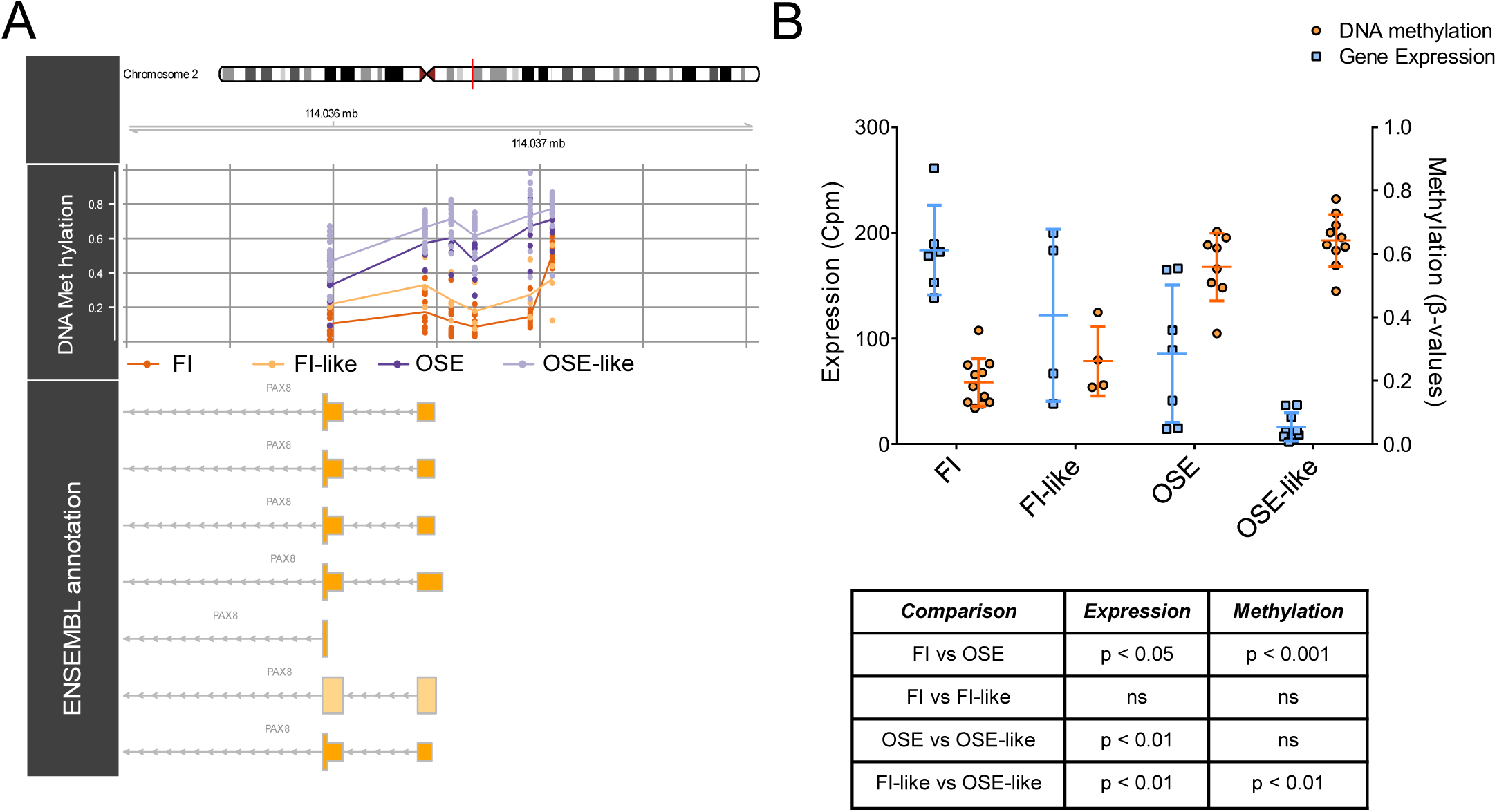
PAX8, a defining marker of HGSOC, is differentially methylated and expressed in FI-like vs. OSE-like tumors. **A.** Graphical representation of the methylation of CpGs in PAX8 promoter across the indicated sample groups **B.** Dotplot depicting the gene expression level of PAX8 by RNAseq (blue bars, left Y-axis) and the DNA methylation level of its promoter by array (orange bars, right Y-axis) in the considered categories. The table summarizes the results of Mann-Whitney U-tests (two-tailed) performed in the indicated comparisons. Data are shown as mean ± standard deviation.

## Discussion

High grade serous ovarian cancer (HGSOC) is a major unmet need in oncology. The lack of suitable human experimental systems that recapitulate the pathogenesis of the tumor is a well-recognized cause of the negligible progress over the last decades^3,4^, with virtually no improvement in patients’ outcome since the introduction of cis-platin as first-line treatment in the 1980s^3,4^. In this regard, the persistent uncertainty regarding the tissue of origin represents a particularly conspicuous hurdle for the elucidation of the molecular pathogenesis as a rational basis for the development of targeted therapies.

In contrast to the previously widely accepted “incessant ovulation” hypothesis^27^, according to which tumors could arise in ovaries as the result of the continuous break/repair of the ovarian surface epithelium, more recent evidence had been increasingly emphasizing the role of the distal tract of the fallopian tube as the most likely tissue of origin of HGSOC, largely as a result of the identification of STIC carrying a core subset of mutations present in both primary and metastatic tumors from the same patient^8^. Concomitantly, however, phylogenetic mutational analysis from different studies posited that a significant fraction of STIC (25%) could actually represent metastatic rather than primary lesions^9^. Along with clear evidence from murine models showing that genetic alterations in both fallopian tube and ovarian surface epithelia can drive tumorigenesis, this contributed to an enduring uncertainty about the relative oncogenic contribution of either epithelium, a situation unlike that of virtually any other solid tumor thanks to the molecular progress of the last decade^28–30^.

Here we tackled and solved this problem by resorting to DNA methylation, given recent evidence pointing to its conservation across normal tissues and their corresponding tumors that warranted its use as a developmental stratifier in multiple contexts^13,14^. We thus derived DNA methylation profiles for both normal (FI and OSE) and tumor (HGSOC and AS) samples. While by analyzing global DNA methylation, an approach adopted by previous more limited studies^7^, we could not distinguish between the two tissues of origin (capturing instead solely the differences existing between normal and tumor samples), we demonstrate that a strategy based on the identification of differentially methylated sites between FI and OSE (OriPrint) allows the robust bipartition of HGSOC, in both cultured cells and whole frozen tissues. Moreover, the identified categories reflect the global variability of HGSOC, thus assigning for the first time a role for the cell of origin in determining the heterogeneity among different tumor samples.

The *a priori* epigenetic stratification in FI-like and OSE-like HGSOC allowed us to interrogate a larger retrospective cohort by the definition of differentially expressed genes between these two categories. Specifically, to interrogate the clinical relevance of our stratification, we resorted to RNAseq profiling of a retrospective cohort of macrodissected FFPE HGSOC samples, employing a shallow 3’-UTR RNAseq approach that allowed us to reduce the library preparation and sequencing costs down to the range of 100 € per sample, thus demonstrating the feasibility of such a stratification for clinical translation. In particular, this approach allows the cost-effective transcriptomic characterization of tumor subtypes through the detection of ~15000 genes, thus avoiding the reduced sensitivity/specificity bound to the restriction to minimal signatures usually employed for classification^31^. By this approach, we found that OSE-like tumors have a negative impact on patients’ prognosis.

BRCA1/2 mutations have previously been linked to increased survival of patients with HGSOC^22^. To check whether the increased survival of FI-like HGSOC affected patients could be due to the co-occurrence of BRCA1/2 mutations, we analysed the occurrence of these mutations on representative subsets of either tumor subtype. We found that BRCA1/2 mutations occur in both types of tumors, thus excluding that these genes are mutated exclusively in the FI-like subtype. Another important aspect linked to DNA Damage Response is the emerging concept of “BRCAness”, according to which tumors can be characterized by mutations/gene inactivation/gene expression patterns whose outcome is similar and closely related to the mutational status of BRCA1/2^32^. While we cannot exclude that our phenotype segregates with the BRCAness phenotype, the comparable distribution of BRCA1/2 mutations among FI-like and OSE.-like HGSOC orients future studies towards the dissection of the role that other alterations in the DDR pathway may potentially contribute to the difference in survival between patients affected by two tumor subtypes.

Finally, in order to gain deeper insight into the molecular features of OSE-like tumors, we analyzed both WGCNA genes and genes that were differentially expressed between FI-like and OSE-like tumors. We identified reduced inflammatory response, higher cell viability, increased cell-to-cell signaling and movement as paradigmatic functions for these tumors. In particular, this is supported by recent evidence showing that the amount of tumor-infiltrating lymphocytes (TILs) correlates with prognosis in patients, and in particular the higher the infiltrate, the better the prognosis^33^. Moreover, the reduced cell death coupled to increased viability and higher cell movement are fully compatible with a more aggressive phenotype in these tumors.

Interestingly, we identified Bromodomain-containing protein 4 (BRD4), a member of the bromodomain and extraterminal (BET) family of chromatin reader proteins, as a positive causal regulator of WGCNA genes in OSE-like tumors. Specifically for HGSOC, BRD4 has been shown to be overexpressed/copy-amplified in a subset of tumors and its amplification has been associated with poor prognosis in patients^34,35^.Moreover, BRD4 overexpression in a OSE cell line was shown to promote tumor transformation, while inhibition of BRD4 in human HGSOC xenografts resulted in reduced tumor volume specifically in BRD4 amplified samples^36^. It will be interesting to test BRD4 inhibition in both subtypes to check whether we could highlight the BRD4 pathway as a specific target for OSE-like tumors.

We further validated these findings by showing that while OSE-like tumors are more enriched in the C1 mesenchymal transcriptional signature, that has been previously shown to correlate with poor prognosis, FI-like tumors are more enriched in the C2 immunoreactive transcriptional signature, a result that is fully compatible with the difference in inflammatory response we scored between the two tumor subtypes and that has been shown to correlate with better prognosis^23^.

Finally, we concentrated on PAX8, a well-known marker of HGSOC and several tumors of Müllerian origin. We showed that PAX8 is differentially expressed and regulated in FI-like vs. OSE-like tumors. Its expression and promoter methylation follow the trend existing in normal tissues, suggesting that it’s expression in tumors could be used as a surrogate lineage tracer, rather than a target for therapy. Nonetheless, evidence shows that the knockdown of PAX8 in HGSOC results in increased apoptosis and reduced proliferation and migration in cancer cell lines^37^. Moreover, the knockdown/overexpression of this gene did not result in tumorigenesis in normal tissues^25,37^, thus suggesting that PAX8 interference could be a potential specific target for HGSOC. Our results build upon this knowledge, allowing to assign PAX8 as a FI-like HGSOC-specific target that could be investigated for improved treatment of this tumor subtype.

In conclusion, our results demonstrate that both fimbrial and ovarian surface epithelium originate HGSOC in humans and establish the feasibility of adopting the OriPRINT-based classification for a rationale stratification of patients. This novel, epigenetically-guided classification has prognostic relevance and illuminates subtype-specific molecular features to define a rationale roadmap towards new therapeutic targets and improved patients’ care.

## Online Methods

### Ethics approval

The study was conducted upon approval of the Ethics Committee of the European Institute of Oncology, Milan following its standard operating procedures (“presa d’atto” 12/3/2014 and 24/7/2017). Fresh tissue samples were obtained upon informed consent from patients undergoing surgery at the Gynecology Division of the European Institute of Oncology. Only tissue samples from patients who have given informed consent to i) the collection of samples for research purposes and their storage into the Biobank of the European Institute of Oncology and ii) the transfer of samples to other research institutions for cancer research purposes have been used in this project. Collected personal data have been pseudonymized, and have been stored and processed in compliance with the applicable data protection legislation, D.Lgs 196/2003 and, since 25 May 2018, Regulation (EU) 2016/679 (General Data Protection Regulation).

### Primary Cells

For all samples, the diagnosis of high grade serous carcinoma was confirmed by pathology review according to the 2014 World Health Organization (WHO) classification (Kurman). High-grade serous epithelial OC cells were derived from peritoneal ascites or from tumor biopsies of patients who had primary, non recurrent OC and had not yet undergone chemotherapy. Primary cells from FI and OSE were derived from non-neoplastic fimbriae and ovaries, respectively, from patients undergoing adnexectomy for non-ovarian gynecological pathologies. The isolation and culture of primary cells were performed as described previously^38,39^. Cells were used at passage 3 to 5.

### Microarray processing and DNA methylation analyses

gDNA from cells was extracted by the DNeasy Blood and Tissue kit (Qiagen) according to manufacturer’s instructions. For each sample 500 ng of genomic DNA were bisulfite converted using the EZ-DNA methylation Gold Kit (Zymo research) according to the kit’s manual, except that the final elution volume was reduced to 12 μl. Per sample, 4 μl of bisulfite converted DNA were used in either the Infinium Human Methylation 450k or the Infinium Methylation EPIC array (both Illumina) procedure according to the vendor’s protocol. Arrays were hybridized according to the manufacturer’s description and scanned on a HiScan system (Illumina). Idat files from IEO cohort and the published datasets were processed using the minfi R package^40^(1.26.0). 450k and EPIC arrays were combined through the combineArrays command (minfi) and preprocessed through SWAN normalization. The three datasets M-values were batch-corrected through ComBat from the SVA package, defining as batch the three datasets and modeling the matrix around the sample types (FI, OSE, HGSOC). To map CpGs to functional elements we used the RnBeads package^41^. To define OriPrint, differential methylation analyses were performed through the limma package^42^ using adj.P.value < 0.05 and logFC > 1 as thresholds. Beta values (logit transformation of M-values) were used for the following analyses. For Pearson’s correlation-based clustering, a distance matrix based on Pearson’s correlation was computed and clustering was performed using the hclust command, using ward.D2 as agglomeration method.

Beta values were imported in Python as anndata object and we used scanpy vs 1.3.1^43^ to plot UMAP and diffusion map for the data. For visualization purposes, we used 20 *n_neighbors*.

Pseudotime analyses were performed with scanpy and the pseudotime origin elements were selected based on their peripheral localization in the multiple dimensionality reduction graphs.

Louvain clustering^20^ was calculated on the first 50 principal components, imposing: 1) minimum number of elements or 2) agglomerating the resulting clusters, with consistent results. HDBScan^21^ was performed on diffusion map’s coordinates, imposing the number of resulting clusters. Gaussian Mixture Model (GMM) clustering was performed with the scikit-learn module, on the first two principal components of variance defined by OriPrint, imposing the number of gaussians and considering the best fitter (log-likelihood) out of 2000 iterations with multiple random initializations.

### Targeted Bisulphite Sequencing

Typically 500 ng of genomic DNA were bisulfite converted using the EZ-DNA methylation Gold Kit (Zymo research) according to the kit’s manual. For PCR amplicons, locus-specific primers (see Table 1) were designed with an in-house tool. PCRs were set-up in 30 μl reactions using 3 μl of 10x Hot Fire Pol Buffer (Solis BioDyne), 4 μl of 10 mM dNTPs (Fisher Scientific), 2.25 μl of 25 mM MgCl2 (Solis BioDyne), 0.6 μl of amplicon specific forward and reverse primer (10 μM each), 0.3 μl of Hot FirePol DNA Polymerase (5 U/μl; Solis Biodyne), 1 μl of bisulfite converted DNA and 18.25 μl of double distilled water. PCRs were run in an ABI Veriti thermo-cycler (Thermo Fisher) using the following program: 95°C for 10 min, then 40 cycles of 95°C for 1 min, 2.5 min of 56°C and 40 sec at 72°C, followed by 7 min of 72°C and hold at 4°C. PCR products were cleaned up using Agencourt AMPure XP Beads (Beckman Coulter). All amplified products were diluted to 4 nM and NGS tags were finalized by a second PCR step (5 cycles) followed by a final clean-up (Agencourt AMPure XP Beads). Finally, all samples (set to 10 nM) were pooled, loaded on an Illumina MiSeq sequencing machine and sequenced for 2x 300 bp paired-end with a MiSeq reagent kit V3 (Illumina) to ca. 10k −20k fold coverage. The raw data was quality checked using FastQC and trimmed for adaptors or low quality bases using the tools cutadapt and Trim Galore!. Paired reads were joined with the FLASh tool. Next, reads were sorted in a two-step procedure by (i) the NGS barcode adaptors to assign Sample ID and (ii) the initial 15 bp to assign amplicon ID. Subsequently, the sorted data was loaded into the BiQAnalyzer HT using the following settings: *analyzed methylation context* was set to “C”, *minimal sequence identity* was set to 0.9 and *minimal conversion rate* was set to 0.95. The filtered high-quality reads were then used for methylation calls of the respective CpGs.

### RNAseq processing and analyses

Total RNA from cells was extracted by the RNeasy Mini Kit (Qiagen) according to manufacturer’s instructions.

H&E-stained slides of all FFPE blocks were assessed for tumor cell content, and the most suitable FFPE block was selected for DNA extraction. For each patient, six tissue shavings per FFPE block, cut at a thickness of 10 μm each, were submitted for RNA extraction. In cases in which the tumor purity in the selected FFPE block was lower than 70%, vital tumor was enriched by macrodissection, scratching unstained slides after the selection of cellular areas by H&E assessment.

Total RNA from FFPE macrodissected tissues was extracted by the RNeasy FFPE kit (Qiagen) according to manufacturer’s instructions on a Qiacube machine (Qiagen).

RNA and further cDNA library quantities were measured using Qubit 2.0 Fluorimetric Assay (Thermo Fisher Scientific) while quality and size were measured by High Sensitivity RNA and DNA screen tapes (Agilent Technologies). Sequencing libraries were constructed starting from 50-100ng of total RNA by optimizations of the QuantSeq 3’ mRNA-Seq Library Prep Kit FWD for Illumina (Lexogen GmbH). DNA libraries were equimolarly pooled at groups of 96 samples and sequenced on a NextSeq 500 high-output, single-end, 75 cycles, v2 Kits (Illumina Inc.).

Sequenced reads were quality-checked using fastQC for read mapping and transcript quantification, we used Salmon (v0.8.1)^44^, we use Hg38 for indexing reference transcripts. Raw counts and transcripts were normalized with TMM using edgeR. All subsequent analyses were conducted using normalized counts. We corrected batch effect within FFPE and fresh sample with SVA^45^ using default parameters. For visualization purposes we used logCPM.

Differential expression analyses were performed with edgeR, setting FDR < 0.05 and logFC > 0.8 to maximize the following analyses (causal network, downstream effect analyses). Gene groups were identified with WGCNA considering among the Module-Trait Relationships (MTRs) those with high correlation with OSE-like tumors.

Survival analysis was performed either considering the 5 years-span overall survival (days elapsed from surgery to death) or the 5 years-span disease-free survival (days elapsed from surgery to relapse’s diagnosis), and calculated using the lifelines package^46^. P value was computed by a standard Logrank Test.

We used a combination of Causal Network Analysis, Downstream Effects Analysis, Upstream Regulator Analysis and Molecule Activity Predictor from Ingenuity Pathway Analysis (QIAGEN Inc. *ingenuity pathway analysis* at https://www.qiagenbioinformatics.com/products/ingenuitypathway-analysis) to identify the predicted functional impact of the genes identified through differential expression or correlation network analysis.

### RT qPCR

1 μg of RNA from each sample was reverse transcribed into cDNA using the Superscript VILO kit (Thermo Fisher) according to manufacturer’s instructions. 200 ng of cDNA were analysed by Taqman (Thermo Fisher) qPCR probing for PAX8 (Hs01015257) GAPDH and ACTB (Hs02758991 and Hs0160665, normalizers) using the SsoAdvanced master mix (Biorad). qPCR was run on a LightCycler 480 (Roche) using the standard amplification protocol for 45 cycles and the average of the Ct of GAPDH and ACTB was used as normalizer. Six independent samples were analyzed in technical triplicates.

## Supporting information

## Author contributions

P.L.R., C.E.V. and G.T. designed experiments and built the intellectual framework, P.L.R. performed experiments including cell culture, nucleic acids extraction and molecular biology assays, C.E.V. and P.L.R. integrated, analysed and interpreted the data, G.G. performed DNA methylation microarray experiments and processed the data, R.L. processed FFPE samples, A.M. and D.C. performed and coordinated the RNAseq processing, A.J. performed targeted bisulphite sequencing experiments and analysed the data, A.V. provided macrodissected FFPE samples, A.V. and A.G. assembled the cohort and provided data on patients, M.L. and U.C. provided biopsy processing and cell culture expertise, provided some primary samples and contributed to experimental discussion, P.L. downloaded and merged published datasets and contributed to discussion on data analysis, V.D. contributed to discussion on data analysis, J.W. contributed to the DNA methylation study design, G.G., A.J., A.V. and D.C. provided the pertinent methods paragraphs, J.W., U.C., N.C., G.V., G.P., and D.C. discussed the data and revised the manuscript, P.L.R., C.E.V. and G.T. wrote the manuscript, P.L.R. and G.T. conceived the project, G.T. supervised the study.

## Acknowledgements

This work was supported by the Associazione Italiana per la Ricerca sul Cancro (AIRC) (IG 2014-2018 to G.T., IG-14622 to U.C.), EPIGEN Flagship Project of the Italian National Research Council (CNR) (to G.T.), Fondazione Italiana per la Ricerca sul Cancro (FIRC) (to P.L.R.), the Italian Ministry of Health (Ricerca Corrente grant to G.T., Ricerca Finalizzata PE-2016-02362551 to U.C.), Fondazione Telethon Core Grant, Armenise-Harvard Foundation Career Development Award, European Research Council (grant agreement 759154, CellKarma), and the Rita-Levi Montalcini program from MIUR (to D.C.), the Fondazione Istituto Europeo di Oncologia-Centro Cardiologico Monzino (to M.L.).

We thank for the support of TIGEM NGS and Bioinformatics Cores. We are grateful to Prof. Pier Paolo Di Fiore who contributed to jump-start the project, to Dr. Cristina Cheroni and Dr. Alessandro Vitriolo for discussion on data analytical tools, to Dr. Giulia Barbagiovanni for fruitful discussion on the work and to Dr. Luca Marelli for the ethical compliance.

