## Supplementary material for "A cell-of-origin epigenetic tracer reveals clinically distinct subtypes of high grade serous ovarian cancer"

### Supplementary Figure 1

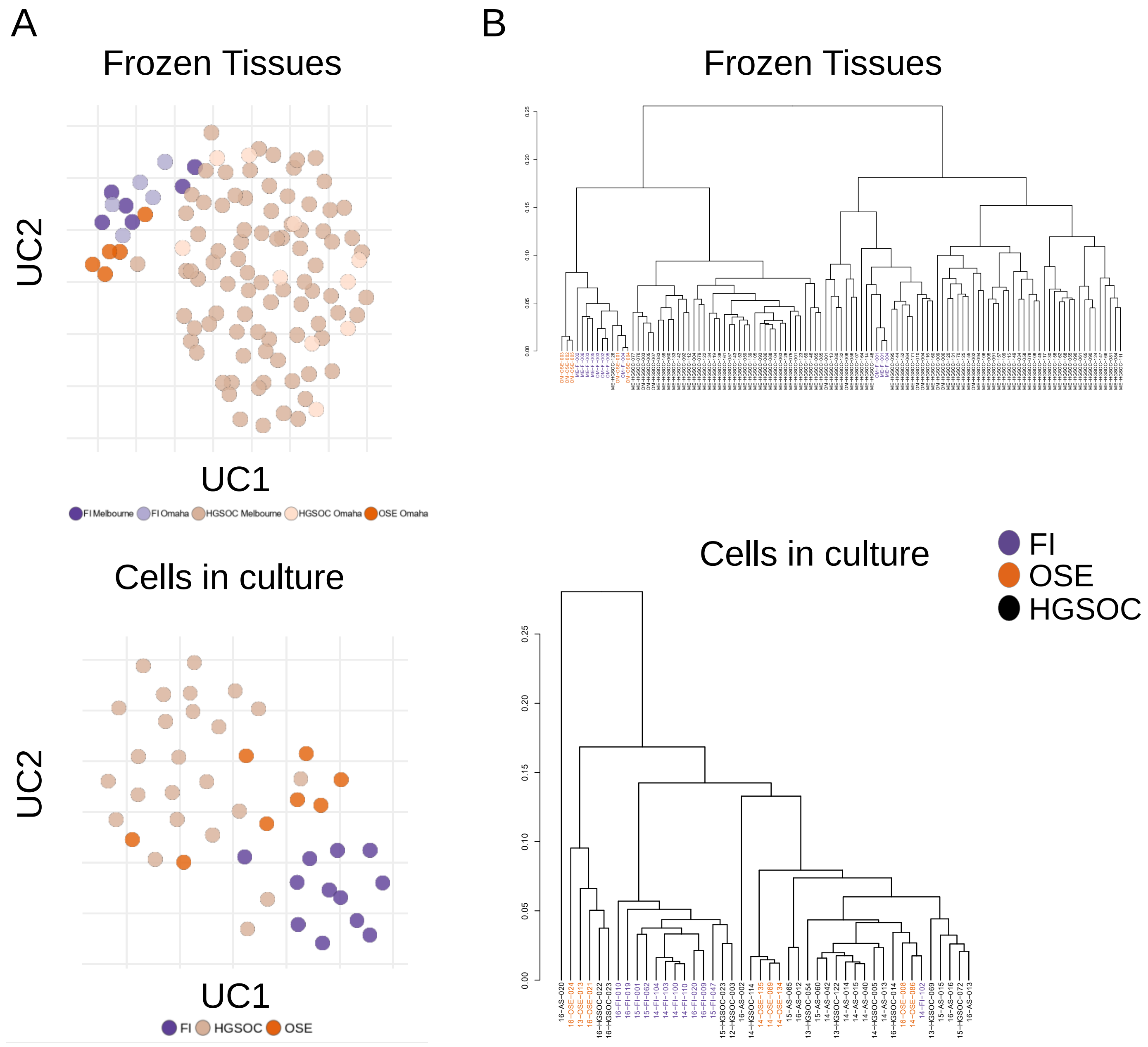


**Supplementary Figure 1. The variance in global DNA methylation does not allow to classify HGSOC according to its cell of origin A.** UMAP plot of the whole methylome of the indicated sample types **B.** Hierarchical clustering performed on the whole methylome of the indicated sample types. Distance= Pearson's correlation.

### Supplementary Figure 2

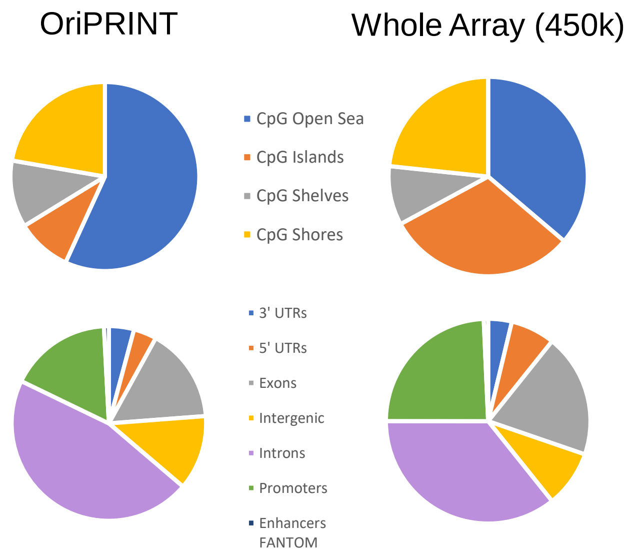


**Supplementary Figure 2. OriPrint CpGs map preferentially to intergenic regions.** Mapping of OriPrint CpGs (left) according to CpG island relatedness (top) or functional regions (bottom). The mapping is compared to the general mapping of the whole Illumina Array (right).

### Supplementary Figure 3

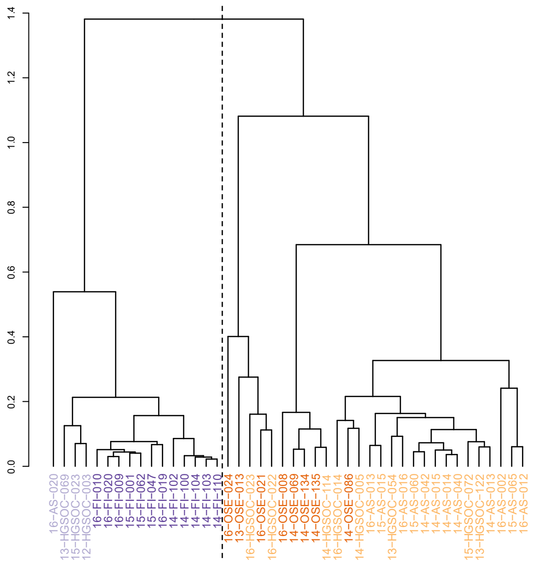


**Supplementary Figure 3. OriPrint allow stratification of tumor samples B.** Hierarchical clustering performed on OriPrint CpGs of the IEO cohort of samples. Distance = Pearson’s correlation.

### Supplementary Figure 4

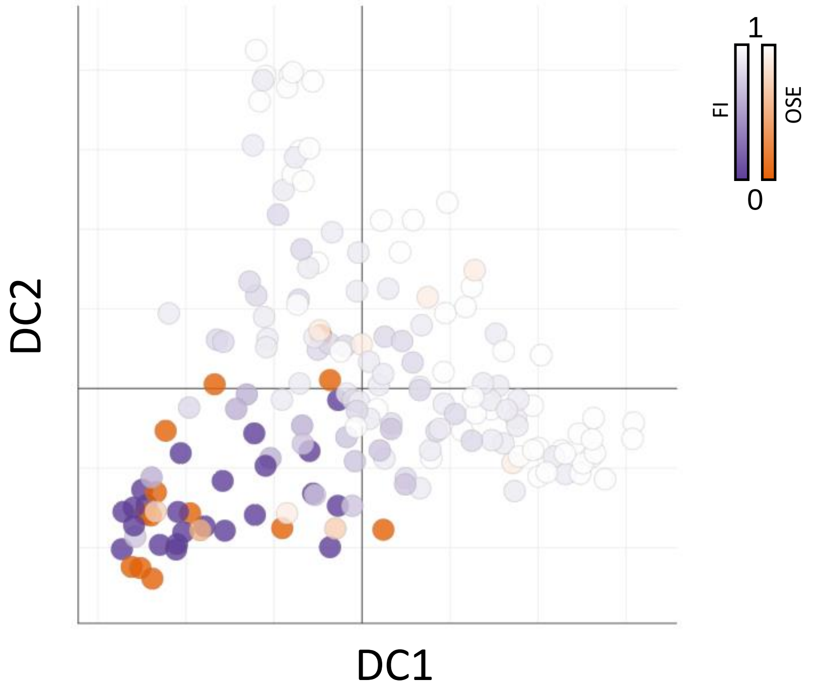


**Supplementary Figure 4. Diffusion pseudotime on global DNA methylome does not allow to derive an evolutionary line from FI and OSE to tumors.** Diffusion map with pseudotime timeline performed on the whole set of CpGs for samples of all cohorts. The origin is situated either in the distal FI (purple origin) or OSE (orange origin) samples.

### Supplementary Figure 5

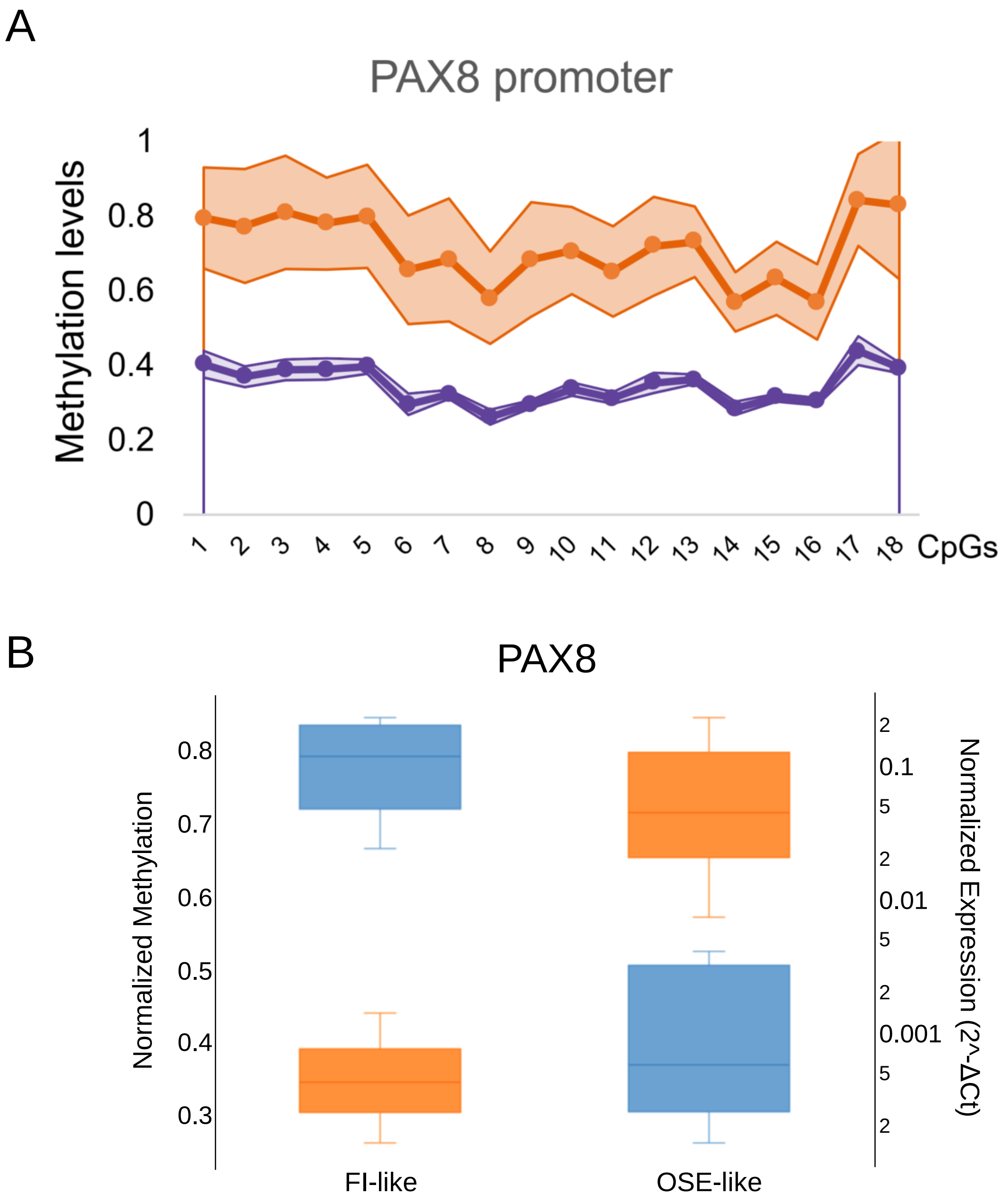


**Supplementary Figure 5. Validation of PAX8 promoter methylation and expression. A:** Methylation levels for the indicated CpG in FI-like (purple) and OSE-like (orange) samples. The data are representative of three independent samples per group. The colored area represents the standard deviation across samples. **B:** Boxplot of PAX8 normalized promoter methylation (orange) and expression (blue) by targeted bisulphite sequencing and qPCR, respectively. Methylation is shown as beta values while expression is shown as 2^(-ΔCt) over the average of GAPDH and ACTB Cts. The values shown are the result of three biological replicates per category (6 samples in total), matched for expression and DNA methylation.
