## Supplementary material for "A cell-of-origin epigenetic tracer reveals clinically distinct subtypes of high grade serous ovarian cancer"

| Sample ID | Hystotype | FIGO Stage | Age at Diagnosis | Weight (kg) | Height (cm) | Date of Diagnosis | Date of Surgery | Date of First Relapse | Date of Last Follow Up | CA125 Pre-Surgery (U/mL) | Ascites (cc) | Metastasis at Lymphnodes | Classification | First Line of Treatment |
|---|---|---|---|---|---|---|---|---|---|---|---|---|---|---|
| FFPE-HGSOC-001 | High Grade Serous Ovarian Cancer | IIIC | 59,72 | 85 | 174 | 2010-05-18 | 2010-01-26 | 2011-03-15 | 2018-01-02 | 481 | 20 | Absent | FI-like | CBDCA-PTX |
| FFPE-HGSOC-002 | High Grade Serous Ovarian Cancer | IIIC | 69,34 | 56 | 150 | 2010-08-16 | 2010-05-16 | 2010-11-05 | 2012-09-01 | 3710 | 0 | Absent | OSE-like | CBDCA-PTX |
| FFPE-HGSOC-003 | High Grade Serous Ovarian Cancer | IIIC | 63,3 | 51 | 150 | 2010-06-01 | 2010-06-08 | 2013-03-06 | 2013-07-10 | 4543 | 20 | Absent | FI-like | CBDCA-PTX |
| FFPE-HGSOC-004 | High Grade Serous Ovarian Cancer | IV | 49,09 | 42 | 148 | 2011-01-27 | 2010-09-03 | 2011-04-01 | 2012-09-08 | 1197 | 0 | Absent | OSE-like | CBDCA-PTX |
| FFPE-HGSOC-005 | High Grade Serous Ovarian Cancer | IIIC | 41,52 | 67 | 160 | 2010-09-02 | 2010-10-12 | 2011-09-21 | 2012-02-15 | 307 | 0 | Absent | FI-like | CBDCA-PTX |
| FFPE-HGSOC-006 | High Grade Serous Ovarian Cancer | IIIC | 66,39 | 58 | 160 | 2011-04-26 | 2011-02-03 | 2012-11-15 | 2015-05-25 | 1530 | 0 | Present | FI-like | CBDCA |
| FFPE-HGSOC-007 | High Grade Serous Ovarian Cancer | IIIC | 83,4 | 48 | 157 | 2011-05-19 | 2011-03-01 | 2011-09-15 | 2016-07-11 | 464 | 0 | Absent | OSE-like | CBDCA-PTX |
| FFPE-HGSOC-008 | High Grade Serous Ovarian Cancer | IIIC | 70,74 | 63 | 150 | 2011-06-28 | 2011-04-05 | 2012-01-13 | 2012-04-29 | 700 | 2000 | Absent | OSE-like | CBDCA-PTX |
| FFPE-HGSOC-009 | High Grade Serous Ovarian Cancer | IIIC | 70,09 | 62 | 154 | 2011-05-20 | 2011-05-20 | 2012-11-07 | 2013-02-05 | 188 | 2000 | Absent | FI-Like | CBDCA-PTX |
| FFPE-HGSOC-010 | High Grade Serous Ovarian Cancer | IV | 58,35 | 56 | 155 | 2011-06-27 | 2011-07-01 | 2012-09-10 | 2013-08-16 | 755 | 50 | Absent | OSE-like | CBDCA-PTX |
| FFPE-HGSOC-011 | High Grade Serous Ovarian Cancer | IIIC | 74,63 | 46 | 156 | 2011-08-25 | 2011-08-25 | 2013-11-15 | 2012-11-12 | 769 | 1400 | Absent | OSE-like | CBDCA-PTX |
| FFPE-HGSOC-012 | High Grade Serous Ovarian Cancer | IIIC | 62,24 | 62 | 157 | 2012-03-05 | 2011-11-25 | 2013-01-23 | 2013-09-16 | 337 | 0 | Absent | OSE-like | CBDCA-PTX |
| FFPE-HGSOC-013 | High Grade Serous Ovarian Cancer | IIIC | 64,44 | 73 | 170 | 2012-04-06 | 2011-12-20 | 2012-07-05 | 2017-03-15 | 2235 | 0 | Absent | OSE-like | CBDCA-PTX |
| FFPE-HGSOC-014 | High Grade Serous Ovarian Cancer | IV | 48,6 | 51 | 155 | 2012-03-19 | 2011-12-20 | 2013-06-17 | 2014-12-30 | 4926 | 0 | Absent | FI-like | CBDCA-PTX |
| FFPE-HGSOC-015 | High Grade Serous Ovarian Cancer | IV | 45,05 | 56 | 169 | 2009-01-08 | 2009-01-08 | 2009-10-08 | 2010-09-14 | 1924 | 4200 | Present | OSE-like | CBDCA-PTX |
| FFPE-HGSOC-016 | High Grade Serous Ovarian Cancer | IIIC | 49,77 | 65 | 162 | 2009-05-12 | 2009-05-12 | 2010-04-15 | 2011-05-15 | 2199 | 1700 | Present | FI-like | CBDCA-PTX |
| FFPE-HGSOC-017 | High Grade Serous Ovarian Cancer | IIIC | 65,00 | 75 | 161 | 2009-11-25 | 2009-11-25 | NaN | 2013-03-24 | 688 | 0 | Present | OSE-like | Other |
| FFPE-HGSOC-018 | High Grade Serous Ovarian Cancer | IIIC | 73,21 | 62 | 150 | 2009-12-15 | 2009-12-15 | 2013-11-27 | 2014-03-14 | 3498 | 2500 | Absent | OSE-like | CBDCA-PTX |
| FFPE-HGSOC-019 | High Grade Serous Ovarian Cancer | IV | 64,87 | 54 | 147 | 2010-01-12 | 2010-01-12 | 2010-02-07 | 2010-02-07 | 5665 | 2000 | Absent | OSE-like | Other |
| FFPE-HGSOC-020 | High Grade Serous Ovarian Cancer | IIIC | 74,48 | 57 | 153 | 2010-02-18 | 2010-02-18 | 2011-06-15 | 2013-01-28 | 1900 | 300 | Absent | OSE-like | CBDCA-PTX |
| FFPE-HGSOC-021 | High Grade Serous Ovarian Cancer | IIIC | 54,78 | 75 | 173 | 2011-02-17 | 2011-02-17 | 2011-12-05 | 2012-05-12 | 2851 | 100 | Present | OSE-like | CBDCA-PTX |
| FFPE-HGSOC-022 | High Grade Serous Ovarian Cancer | IV | 45,83 | 67 | 160 | 2011-03-17 | 2011-03-17 | 2012-04-17 | 2012-07-15 | 517 | 2800 | Present | OSE-like | CBDCA-PTX |
| FFPE-HGSOC-023 | High Grade Serous Ovarian Cancer | IIIC | 72,08 | 54 | 165 | 2011-06-07 | 2011-06-07 | 2012-04-15 | 2013-02-15 | 535 | 1400 | Absent | OSE-like | CBDCA-PTX |
| FFPE-HGSOC-024 | High Grade Serous Ovarian Cancer | IV | 62,54 | 93 | 168 | 2012-03-14 | 2012-03-14 | 2012-05-13 | 2012-05-13 | 1225 | 400 | Present | OSE-like | CBDCA-PTX |
| FFPE-HGSOC-025 | High Grade Serous Ovarian Cancer | IIIC | 62,74 | 68 | 162 | 2012-04-11 | 2012-04-11 | 2012-10-11 | 2013-01-17 | 2689 | 5000 | Absent | FI-like | CBDCA-PTX |
| FFPE-HGSOC-026 | High Grade Serous Ovarian Cancer | IIIC | 56,49 | 44 | 150 | 2012-04-20 | 2012-04-20 | 2012-08-22 | 2013-02-11 | 804 | 1500 | Absent | FI-like | CBDCA-PTX |
| FFPE-HGSOC-027 | High Grade Serous Ovarian Cancer | IV | 52,27 | 56 | 154 | 2009-01-09 | 2009-01-09 | 2010-01-11 | 2011-12-22 | 2753 | 1000 | Present | OSE-like | CBDCA-PTX |
| FFPE-HGSOC-028 | High Grade Serous Ovarian Cancer | IV | 63,92 | 66 | 158 | 2009-09-23 | 2009-09-23 | 2011-01-08 | 2012-05-01 | 1852 | 0 | Absent | OSE-like | CBDCA-PTX |
| FFPE-HGSOC-029 | High Grade Serous Ovarian Cancer | IV | 67,46 | 52 | 162 | 2009-02-10 | 2009-02-10 | 2009-12-17 | 2011-02-15 | 1749 | 0 | Present | FI-like | CBDCA-PTX |
| FFPE-HGSOC-030 | High Grade Serous Ovarian Cancer | IIIC | 38,27 | 59 | 160 | 2009-02-11 | 2009-02-11 | 2010-04-14 | 2011-05-14 | 2154 | 2000 | Absent | FI-like | CBDCA-PTX |
| FFPE-HGSOC-031 | High Grade Serous Ovarian Cancer | IV | 47,34 | 56 | 170 | 2009-02-13 | 2009-02-13 | 2010-04-01 | 2012-01-15 | 2685 | 500 | Present | OSE-like | CBDCA-PTX |
| FFPE-HGSOC-032 | High Grade Serous Ovarian Cancer | IIIC | 54,24 | 64 | 165 | 2009-02-17 | 2009-02-17 | 2009-11-09 | 2011-09-21 | 740 | 5200 | Present | OSE-like | CBDCA |
| FFPE-HGSOC-033 | High Grade Serous Ovarian Cancer | IV | 44,98 | 50 | 164 | 2009-02-19 | 2009-02-19 | 2010-04-12 | 2011-04-11 | 4176 | 2000 | Absent | OSE-like | CBDCA-PTX |
| FFPE-HGSOC-034 | High Grade Serous Ovarian Cancer | IV | 51,07 | 45 | 153 | 2009-03-20 | 2009-03-20 | 2009-08-12 | 2010-05-10 | 1185 | 1500 | Present | OSE-like | CBDCA |
| FFPE-HGSOC-035 | High Grade Serous Ovarian Cancer | IV | 60,12 | 50 | 162 | 2009-03-25 | 2009-03-25 | 2010-04-15 | 2010-04-15 | 3517 | 300 | Absent | OSE-like | CBDCA-PTX |
| FFPE-HGSOC-036 | High Grade Serous Ovarian Cancer | IIIC | 50,00 | 78 | 160 | 2009-03-31 | 2009-03-31 | 2011-03-15 | 2014-02-11 | 3769 | 3000 | Absent | FI-like | CBDCA-PTX |
| FFPE-HGSOC-037 | High Grade Serous Ovarian Cancer | IIIC | 62,71 | 56 | 153 | 2009-06-24 | 2009-06-24 | 2010-07-01 | 2012-09-29 | 124 | 0 | Present | OSE-like | CBDCA-PTX |
| FFPE-HGSOC-038 | High Grade Serous Ovarian Cancer | IIIC | 70,90 | 54 | 159 | 2009-07-17 | 2009-07-17 | 2010-08-02 | 2010-11-27 | 166 | 500 | Present | OSE-like | CBDCA-PTX |
| FFPE-HGSOC-039 | High Grade Serous Ovarian Cancer | IV | 58,74 | 64 | 165 | 2009-07-21 | 2009-07-21 | 2010-08-13 | 2012-10-15 | 553 | 500 | Present | OSE-like | CBDCA-PTX |
| FFPE-HGSOC-040 | High Grade Serous Ovarian Cancer | IV | 47,54 | 58 | 152 | 2009-08-12 | 2009-08-12 | 2011-05-15 | 2011-09-16 | 4116 | 3000 | Present | OSE-like | CBDCA-PTX |
| FFPE-HGSOC-041 | High Grade Serous Ovarian Cancer | IIIC | 67,34 | 73 | 164 | 2009-08-25 | 2009-08-25 | 2010-07-30 | 2014-01-15 | 1328 | 2500 | Absent | OSE-like | CBDCA-PTX |
| FFPE-HGSOC-042 | High Grade Serous Ovarian Cancer | IV | 60,55 | 60 | 159 | 2009-09-15 | 2009-09-15 | 2010-05-15 | 2010-07-23 | 17412 | 3000 | Present | OSE-like | CBDCA-PTX |
| FFPE-HGSOC-043 | High Grade Serous Ovarian Cancer | IV | 62,57 | 65 | 158 | 2009-09-29 | 2009-09-29 | 2010-09-01 | 2013-11-13 | 4265 | 0 | Absent | OSE-like | CBDCA-PTX |
| FFPE-HGSOC-044 | High Grade Serous Ovarian Cancer | IIIC | 49,83 | 80 | 161 | 2009-10-22 | 2009-10-22 | 2010-09-02 | 2012-10-04 | 459 | 4000 | Absent | OSE-like | CBDCA-PTX |
| FFPE-HGSOC-045 | High Grade Serous Ovarian Cancer | IIIC | 56,24 | 54 | 168 | 2009-10-27 | 2009-10-27 | 2011-04-08 | 2012-04-29 | 5976 | 50 | Present | OSE-like | CBDCA-PTX |
| FFPE-HGSOC-046 | High Grade Serous Ovarian Cancer | IIIC | 47,15 | 61 | 165 | 2009-11-10 | 2009-11-10 | 2010-06-22 | 2011-07-28 | 1400 | 30 | Present | OSE-like | CBDCA-PTX |
| FFPE-HGSOC-047 | High Grade Serous Ovarian Cancer | IIIC | 52,54 | 60 | 179 | 2010-01-05 | 2010-01-05 | 2010-06-20 | 2010-08-29 | 130 | 0 | Absent | FI-like | CBDCA-PTX |
| FFPE-HGSOC-048 | High Grade Serous Ovarian Cancer | IV | 50,52 | 69 | 169 | 2010-01-27 | 2010-01-27 | 2010-12-14 | 2011-04-26 | 3569 | 3000 | Present | OSE-like | CBDCA-PTX |
| FFPE-HGSOC-049 | High Grade Serous Ovarian Cancer | IIIC | 43,40 | 61 | 170 | 2010-02-10 | 2010-02-10 | 2010-11-18 | 2011-11-06 | 4247 | 500 | Present | OSE-like | CBDCA-PTX |
| FFPE-HGSOC-050 | High Grade Serous Ovarian Cancer | IIIC | 43,27 | 89 | 170 | 2010-02-17 | 2010-02-17 | 2011-03-01 | 2013-08-19 | 1733 | 5000 | Present | OSE-like | CBDCA-PTX |
| FFPE-HGSOC-051 | High Grade Serous Ovarian Cancer | IIIC | 62,80 | 62 | 160 | 2010-02-24 | 2010-02-24 | 2012-07-26 | 2012-07-26 | 835 | 1000 | Present | FI-like | CBDCA-PTX |
| FFPE-HGSOC-052 | High Grade Serous Ovarian Cancer | IIIC | 51,64 | 80 | 163 | 2010-03-03 | 2010-03-03 | 2010-07-23 | 2010-09-09 | 294 | 4000 | Present | OSE-like | CBDCA-PTX |
| FFPE-HGSOC-053 | High Grade Serous Ovarian Cancer | IV | 63,87 | 91 | 167 | 2010-03-03 | 2010-03-03 | 2011-05-03 | 2013-07-26 | 1273 | 2700 | Absent | OSE-like | CBDCA-PTX |
| FFPE-HGSOC-054 | High Grade Serous Ovarian Cancer | IIIC | 69,72 | 85 | 169 | 2010-03-19 | 2010-03-19 | 2011-03-01 | 2011-03-01 | 1620 | 0 | Present | OSE-like | CBDCA-PTX |
| FFPE-HGSOC-055 | High Grade Serous Ovarian Cancer | IIIC | 42,00 | 69 | 172 | 2010-04-16 | 2010-04-16 | 2010-07-30 | 2011-03-01 | 206 | 200 | Absent | OSE-like | CBDCA-PTX |
| FFPE-HGSOC-056 | High Grade Serous Ovarian Cancer | IV | 62,27 | 45 | 165 | 2010-04-27 | 2010-04-27 | NaN | 2012-02-18 | 15835 | 1500 | Present | OSE-like | Other |
| FFPE-HGSOC-057 | High Grade Serous Ovarian Cancer | IV | 62,86 | 52 | 165 | 2010-05-13 | 2010-05-13 | 2011-03-24 | 2012-06-23 | 3639 | 500 | Present | OSE-like | CBDCA-PTX |
| FFPE-HGSOC-058 | High Grade Serous Ovarian Cancer | IIIC | 55,14 | 64 | 160 | 2010-05-19 | 2010-05-19 | 2012-01-05 | 2012-08-11 | 6046 | 3500 | Present | OSE-like | CBDCA-PTX |
| FFPE-HGSOC-059 | High Grade Serous Ovarian Cancer | IIIC | 64,70 | 60 | 155 | 2010-06-18 | 2010-06-18 | 2011-05-30 | 2013-12-26 | 1343 | 200 | Absent | OSE-like | CBDCA-PTX |
| FFPE-HGSOC-060 | High Grade Serous Ovarian Cancer | IIIC | 52,06 | 95 | 168 | 2010-06-23 | 2010-06-23 | 2011-10-27 | 2014-05-21 | 1621 | 1000 | Absent | FI-like | CBDCA-PTX |
| FFPE-HGSOC-061 | High Grade Serous Ovarian Cancer | IIIC | 66,03 | 66 | 158 | 2010-07-06 | 2010-07-06 | 2011-11-02 | 2013-05-04 | 600 | 6000 | Present | FI-like | CBDCA-PTX |
| FFPE-HGSOC-062 | High Grade Serous Ovarian Cancer | IIIC | 63,72 | 51 | 165 | 2010-07-28 | 2010-07-28 | 2011-01-24 | 2013-04-19 | 32884 | 0 | Present | OSE-like | CBDCA-PTX |
| FFPE-HGSOC-063 | High Grade Serous Ovarian Cancer | IIIC | 62,86 | 80 | 155 | 2010-08-03 | 2010-08-03 | NaN | 2011-05-01 | 7084 | 100 | Present | OSE-like | Other |
| FFPE-HGSOC-064 | High Grade Serous Ovarian Cancer | IV | 64,88 | 45 | 160 | 2010-08-30 | 2010-08-30 | 2011-07-06 | 2012-11-22 | 1420 | 1800 | Present | FI-like | CBDCA-PTX |
| FFPE-HGSOC-065 | High Grade Serous Ovarian Cancer | IIIC | 51,49 | 54 | 177 | 2010-09-20 | 2010-09-20 | 2011-09-29 | 2012-07-01 | 777 | 2400 | Absent | OSE-like | CBDCA-PTX |
| FFPE-HGSOC-066 | High Grade Serous Ovarian Cancer | IV | 44,19 | 95 | 169 | 2010-10-06 | 2010-10-06 | 2011-06-21 | 2012-12-16 | 483 | 1200 | Present | OSE-like | CBDCA |
| FFPE-HGSOC-067 | High Grade Serous Ovarian Cancer | IIIC | 62,87 | 92 | 161 | 2010-11-22 | 2010-11-22 | NaN | 2010-11-25 | 1200 | 2200 | Present | OSE-like | Other |
| FFPE-HGSOC-068 | High Grade Serous Ovarian Cancer | IIIC | 71,32 | 53 | 146 | 2010-11-24 | 2010-11-24 | 2012-03-21 | 2012-03-21 | 1806 | 2500 | Present | OSE-like | CBDCA |
| FFPE-HGSOC-069 | High Grade Serous Ovarian Cancer | IIIC | 58,79 | 75 | 161 | 2010-11-30 | 2010-11-30 | 2011-11-10 | 2013-07-23 | 1968 | 2000 | Absent | FI-like | CBDCA-PTX |
| FFPE-HGSOC-070 | High Grade Serous Ovarian Cancer | IV | 51,47 | 48 | 157 | 2011-01-18 | 2011-01-18 | 2011-09-02 | 2012-11-21 | 7277 | 3000 | Present | OSE-like | CBDCA-PTX |
| FFPE-HGSOC-071 | High Grade Serous Ovarian Cancer | IV | 42,38 | 71 | 171 | 2009-03-18 | 2009-03-18 | 2011-01-18 | 2018-10-18 | 11972 | 7000 | Present | FI-like | CBDCA-PTX |
| FFPE-HGSOC-072 | High Grade Serous Ovarian Cancer | IIIC | 71,60 | 60 | 155 | 2009-04-17 | 2009-04-17 | 2010-09-15 | 2015-02-17 | 2313 | 100 | Present | OSE-like | CBDCA-PTX |
| FFPE-HGSOC-073 | High Grade Serous Ovarian Cancer | IIIC | 70,71 | 51 | 161 | 2009-06-30 | 2009-06-30 | 2011-06-29 | 2017-03-15 | 55 | 0 | Present | FI-like | CBDCA-PTX |
| FFPE-HGSOC-074 | High Grade Serous Ovarian Cancer | IIIC | 57,72 | 71 | 153 | 2009-08-26 | 2009-08-26 | 2010-03-15 | 2017-03-15 | 143 | 3500 | Present | FI-like | CBDCA-PTX |
| FFPE-HGSOC-075 | High Grade Serous Ovarian Cancer | IIIC | 62,56 | 72 | 160 | 2009-09-02 | 2009-09-02 | 2011-09-02 | 2017-03-15 | 115 | 0 | Present | FI-like | CBDCA-PTX |
| FFPE-HGSOC-076 | High Grade Serous Ovarian Cancer | IIIC | 42,23 | 59 | 160 | 2009-12-23 | 2009-12-23 | 2011-10-18 | 2018-03-02 | 389 | 0 | Present | OSE-like | CBDCA-PTX |
| FFPE-HGSOC-077 | High Grade Serous Ovarian Cancer | IIIC | 62,81 | 57 | 165 | 2010-10-19 | 2010-10-19 | 2011-06-01 | 2017-03-15 | 462 | 3500 | Absent | FI-like | CBDCA-PTX |
| FFPE-HGSOC-078 | High Grade Serous Ovarian Cancer | IIIC | 72,25 | 80 | 168 | 2010-11-16 | 2010-11-16 | 2011-05-15 | 2017-03-15 | 892 | 5000 | Absent | OSE-like | CBDCA-PTX |
| FFPE-HGSOC-079 | High Grade Serous Ovarian Cancer | IV | 43,47 | 80 | 167 | 2011-02-14 | 2011-02-14 | 2012-04-10 | 2014-11-16 | 651 | 1500 | Present | OSE-like | CBDCA-PTX |
| FFPE-HGSOC-080 | High Grade Serous Ovarian Cancer | IIIC | 45,57 | 62 | 165 | 2011-03-29 | 2011-03-29 | 2013-06-28 | 2014-12-20 | 411 | 100 | Present | FI-like | CBDCA-PTX |
| FFPE-HGSOC-081 | High Grade Serous Ovarian Cancer | IIIC | 70,91 | 100 | 165 | 2011-04-13 | 2011-04-13 | 2012-03-15 | 2013-12-31 | 553 | 1000 | Present | FI-like | CBDCA-PTX |
| FFPE-HGSOC-082 | High Grade Serous Ovarian Cancer | IIIC | 79,73 | 63 | 163 | 2011-04-15 | 2011-04-15 | 2011-09-20 | 2012-09-09 | 20 | 0 | Absent | OSE-like | CBDCA |
| FFPE-HGSOC-083 | High Grade Serous Ovarian Cancer | IIIC | 52,28 | 69 | 160 | 2011-04-28 | 2011-04-28 | 2012-08-23 | 2015-02-07 | 2687 | 700 | Absent | OSE-like | CBDCA-PTX |
| FFPE-HGSOC-084 | High Grade Serous Ovarian Cancer | IIIC | 46,33 | 54 | 155 | 2011-06-15 | 2011-06-15 | 2011-09-19 | 2011-09-19 | 745 | 0 | Present | FI-like | CBDCA-PTX |
| FFPE-HGSOC-085 | High Grade Serous Ovarian Cancer | IIIC | 73,82 | 60 | 153 | 2011-07-04 | 2011-07-04 | 2012-08-13 | 2015-02-26 | 3276 | 350 | Absent | OSE-like | CBDCA |
| FFPE-HGSOC-086 | High Grade Serous Ovarian Cancer | IIIC | 62,34 | 155 | 155 | 2011-08-25 | 2011-08-25 | 2013-01-07 | 2014-08-26 | 2696 | 1000 | Present | OSE-like | CBDCA-PTX |

| | | | | | | | | | | | | | | |
|---|---|---|---|---|---|---|---|---|---|---|---|---|---|---|
| FFPE-HGSOC-087 | High Grade Serous Ovarian Cancer | IV | 75,12 | 58 | 155 | 2011-09-05 | 2011-09-05 | 2012-08-01 | 2012-11-21 | 545 | 100 | Present | OSE-like | CBDCA-PTX |
| FFPE-HGSOC-088 | High Grade Serous Ovarian Cancer | IV | 67,39 | 63 | 164 | 2011-11-10 | 2011-11-10 | 2013-09-11 | 2018-10-01 | 319 | 50 | Absent | OSE-like | CBDCA-PTX |
| FFPE-HGSOC-089 | High Grade Serous Ovarian Cancer | IIIC | 67,04 | 72 | 163 | 2011-11-30 | 2011-11-30 | 2012-09-20 | 2013-02-15 | 686 | 500 | Present | OSE-like | CBDCA-PTX |
| FFPE-HGSOC-090 | High Grade Serous Ovarian Cancer | IIIC | 77,46 | 80 | 165 | 2012-01-09 | 2012-01-09 | 2012-07-23 | 2013-12-04 | 2073 | 1800 | Absent | OSE-like | CBDCA |
| FFPE-HGSOC-091 | High Grade Serous Ovarian Cancer | IIIC | 74,05 | 76 | 160 | 2012-02-20 | 2012-02-20 | 2012-03-22 | 2012-03-22 | 5392 | 7000 | Absent | OSE-like | Other |
| FFPE-HGSOC-092 | High Grade Serous Ovarian Cancer | IIIC | 74,37 | 55 | 162 | 2012-07-23 | 2012-07-23 | 2012-12-13 | 2017-03-21 | 4584 | 3800 | Absent | FI-like | CBDCA-PTX |
| FFPE-HGSOC-093 | High Grade Serous Ovarian Cancer | IV | 75,59 | 63 | 170 | 2012-07-25 | 2012-07-25 | 2012-08-12 | 2012-08-12 | 42 | 0 | Absent | FI-like | CBDCA-PTX |
| FFPE-HGSOC-094 | High Grade Serous Ovarian Cancer | IIIC | 74,13 | 74 | 170 | 2012-08-14 | 2012-08-14 | 2013-03-01 | 2014-03-31 | 621 | 2800 | Present | FI-like | CBDCA |
| FFPE-HGSOC-095 | High Grade Serous Ovarian Cancer | IV | 58,18 | 60 | 168 | 2012-10-19 | 2012-10-19 | 2013-03-18 | 2013-10-21 | 21125 | 6000 | Present | FI-like | CBDCA-PTX |
| FFPE-HGSOC-096 | High Grade Serous Ovarian Cancer | IIIC | 42,69 | 60 | 160 | 2012-11-14 | 2012-11-14 | 2013-10-15 | 2017-04-11 | 45 | 0 | Absent | OSE-like | CBDCA-PTX |
| FFPE-HGSOC-097 | High Grade Serous Ovarian Cancer | IIIC | 57,75 | 50 | 155 | 2010-02-02 | 2010-02-02 | 2011-08-23 | 2013-05-02 | 251 | 30 | Absent | FI-like | CBDCA-PTX |
| FFPE-HGSOC-098 | High Grade Serous Ovarian Cancer | IV | 51,15 | 83 | 158 | 2010-02-23 | 2010-02-23 | 2011-11-29 | 2018-10-16 | 5109 | 0 | Present | FI-like | CBDCA-PTX |
| FFPE-HGSOC-099 | High Grade Serous Ovarian Cancer | IIIC | 52,55 | 73 | 165 | 2010-02-25 | 2010-02-25 | 2011-11-04 | 2017-03-15 | 2968 | 6200 | Absent | FI-like | CBDCA-PTX |
| FFPE-HGSOC-100 | High Grade Serous Ovarian Cancer | IIIC | 51,36 | 70 | 160 | 2010-03-17 | 2010-03-17 | 2011-09-21 | 2016-12-17 | 533 | 700 | Absent | OSE-like | CBDCA-PTX |
| FFPE-HGSOC-101 | High Grade Serous Ovarian Cancer | IV | 45,34 | 48 | 158 | 2010-03-30 | 2010-03-30 | 2011-04-15 | 2013-08-12 | 7375 | 3000 | Present | FI-like | CBDCA-PTX |
| FFPE-HGSOC-102 | High Grade Serous Ovarian Cancer | IIIC | 58,09 | 69 | 170 | 2010-05-28 | 2010-05-28 | 2013-02-23 | 2017-02-09 | 523 | 0 | Absent | FI-like | CBDCA-PTX |
| FFPE-HGSOC-103 | High Grade Serous Ovarian Cancer | IIIC | 74,52 | 56 | 162 | 2010-06-23 | 2010-06-23 | 2011-10-06 | 2017-06-03 | 6831 | 500 | Present | FI-like | CBDCA-PTX |
| FFPE-HGSOC-104 | High Grade Serous Ovarian Cancer | IIIC | 72,72 | 83 | 160 | 2010-07-16 | 2010-07-16 | 2014-03-15 | 2017-03-15 | 5467 | 0 | Absent | FI-like | Other |
| FFPE-HGSOC-105 | High Grade Serous Ovarian Cancer | IIIC | 56,79 | 68 | 155 | 2010-07-19 | 2010-07-19 | 2013-07-08 | 2017-02-15 | 4472 | 1500 | Absent | FI-like | CBDCA-PTX |
| FFPE-HGSOC-106 | High Grade Serous Ovarian Cancer | IIIC | 67,39 | 67 | 168 | 2010-07-23 | 2010-07-23 | 2013-10-28 | 2015-03-22 | 2882 | 11000 | Absent | OSE-like | CBDCA-PTX |
| FFPE-HGSOC-107 | High Grade Serous Ovarian Cancer | IV | 73,35 | 52 | 170 | 2010-07-30 | 2010-07-30 | 2012-02-15 | 2017-03-15 | 886 | 2700 | Absent | FI-like | CBDCA-PTX |
| FFPE-HGSOC-108 | High Grade Serous Ovarian Cancer | IV | 54,51 | 56 | 164 | 2010-08-06 | 2010-08-06 | 2013-05-03 | 2017-03-15 | 4906 | 0 | Present | OSE-like | CBDCA |
| FFPE-HGSOC-109 | High Grade Serous Ovarian Cancer | IIIC | 67,37 | 72 | 160 | 2010-09-28 | 2010-09-28 | 2012-06-07 | 2017-03-15 | 745 | 8000 | Present | OSE-like | CBDCA-PTX |
| FFPE-HGSOC-110 | High Grade Serous Ovarian Cancer | IIIC | 48,09 | 58 | 164 | 2010-10-22 | 2010-10-22 | 2012-04-19 | 2018-10-17 | 295 | 0 | Present | OSE-like | CBDCA-PTX |
| FFPE-HGSOC-111 | High Grade Serous Ovarian Cancer | IIIC | 59,80 | 64 | 162 | 2010-11-02 | 2010-11-02 | 2012-09-05 | 2017-01-25 | 391 | 0 | Present | FI-like | CBDCA-PTX |
| FFPE-HGSOC-112 | High Grade Serous Ovarian Cancer | IIIC | 65,45 | 70 | 170 | 2010-11-29 | 2010-11-29 | 2012-03-03 | 2013-02-27 | 413 | 100 | Absent | FI-like | CBDCA-PTX |
| FFPE-HGSOC-113 | High Grade Serous Ovarian Cancer | IIIB | 63,68 | 64 | 166 | 2010-12-13 | 2010-12-13 | 2011-10-28 | 2014-08-21 | 1866 | 7000 | Absent | FI-like | CBDCA-PTX |
| FFPE-HGSOC-114 | High Grade Serous Ovarian Cancer | IIIC | 64,94 | 72 | 168 | 2010-12-21 | 2010-12-21 | 2012-06-15 | 2015-09-19 | 3454 | 1500 | Absent | FI-like | CBDCA-PTX |
| FFPE-HGSOC-115 | High Grade Serous Ovarian Cancer | IIIC | 65,89 | 49 | 165 | 2010-12-23 | 2010-12-23 | 2011-11-14 | 2018-07-06 | 1479 | 0 | Present | OSE-like | CBDCA-PTX |
| FFPE-HGSOC-116 | High Grade Serous Ovarian Cancer | IIIC | 66,82 | 59 | 150 | 2011-02-16 | 2011-02-16 | 2013-01-10 | 2016-06-15 | 794 | 300 | Present | OSE-like | CBDCA-PTX |
| FFPE-HGSOC-117 | High Grade Serous Ovarian Cancer | IIIC | 54,99 | 60 | 160 | 2011-03-01 | 2011-03-01 | 2012-06-13 | 2017-03-15 | 1479 | 300 | Present | FI-like | CBDCA-PTX |
| FFPE-HGSOC-118 | High Grade Serous Ovarian Cancer | IIIC | 51,09 | 61 | 172 | 2011-03-15 | 2011-03-15 | 2014-05-12 | 2018-08-22 | 117 | 0 | Absent | FI-like | CBDCA-PTX |
| FFPE-HGSOC-119 | High Grade Serous Ovarian Cancer | IIIC | 64,86 | 67 | 173 | 2011-03-16 | 2011-03-16 | 2012-01-04 | 2018-04-18 | 1245 | 300 | Absent | FI-like | CBDCA-PTX |
| FFPE-HGSOC-120 | High Grade Serous Ovarian Cancer | IIIC | 50,05 | 55 | 168 | 2011-03-29 | 2011-03-29 | 2013-12-15 | 2017-03-15 | 1325 | 800 | Absent | FI-like | CBDCA-PTX |
| FFPE-HGSOC-121 | High Grade Serous Ovarian Cancer | IV | 58,66 | 52 | 175 | 2011-03-31 | 2011-03-31 | 2013-01-10 | 2018-10-11 | 70 | 0 | Absent | FI-like | CBDCA-PTX |
| FFPE-HGSOC-122 | High Grade Serous Ovarian Cancer | IIIC | 69,25 | 68 | 160 | 2011-04-06 | 2011-04-06 | 2012-10-23 | 2014-11-01 | 781 | 0 | Present | FI-like | CBDCA-PTX |
| FFPE-HGSOC-123 | High Grade Serous Ovarian Cancer | IV | 59,96 | 122 | 167 | 2011-04-18 | 2011-04-18 | 2014-04-09 | 2018-10-09 | 957 | 7000 | Absent | FI-like | CBDCA-PTX |
| FFPE-HGSOC-124 | High Grade Serous Ovarian Cancer | IIIC | 47,84 | 54 | 165 | 2011-05-30 | 2011-05-30 | 2014-04-03 | 2016-02-03 | 58 | 8000 | Absent | OSE-like | CBDCA-PTX |
| FFPE-HGSOC-125 | High Grade Serous Ovarian Cancer | IIIC | 44,10 | 50 | 154 | 2011-06-01 | 2011-06-01 | 2012-08-30 | 2016-11-21 | 50 | 0 | Present | OSE-like | CBDCA-PTX |
| FFPE-HGSOC-126 | High Grade Serous Ovarian Cancer | IIIC | 62,87 | 84 | 168 | 2011-07-08 | 2011-07-08 | 2013-06-05 | 2015-05-08 | 1845 | 2000 | Absent | OSE-like | CBDCA-PTX |
| FFPE-HGSOC-127 | High Grade Serous Ovarian Cancer | IV | 55,76 | 153 | 153 | 2011-07-25 | 2011-07-25 | 2013-02-13 | 2015-02-10 | 314 | 1300 | Absent | OSE-like | CBDCA-PTX |
| FFPE-HGSOC-128 | High Grade Serous Ovarian Cancer | IIIC | 62,84 | 160 | 160 | 2011-08-26 | 2011-08-26 | 2013-10-30 | 2018-06-25 | 4163 | 700 | Absent | OSE-like | CBDCA-PTX |
| FFPE-HGSOC-129 | High Grade Serous Ovarian Cancer | IIIC | 52,53 | 56 | 155 | 2011-09-21 | 2011-09-21 | 2014-03-26 | 2015-09-01 | 3324 | 5400 | Absent | OSE-like | CBDCA-PTX |
| FFPE-HGSOC-130 | High Grade Serous Ovarian Cancer | IIIC | 48,31 | 58 | 160 | 2011-11-24 | 2011-11-24 | 2013-07-02 | 2015-05-30 | 4404,7 | 1500 | Present | OSE-like | CBDCA-PTX |
| FFPE-HGSOC-131 | High Grade Serous Ovarian Cancer | IIIC | 82,07 | 78 | 170 | 2011-12-02 | 2011-12-02 | 2013-07-02 | 2015-01-24 | 30 | 0 | Present | OSE-like | CBDCA |
| FFPE-HGSOC-132 | High Grade Serous Ovarian Cancer | IIIC | 54,58 | 80 | 160 | 2011-12-16 | 2011-12-16 | 2013-09-06 | 2016-01-15 | 637 | 8000 | Present | FI-like | CBDCA-PTX |
| FFPE-HGSOC-133 | High Grade Serous Ovarian Cancer | IV | 49,08 | 41 | 153 | 2012-02-06 | 2012-02-06 | 2013-10-09 | 2017-03-15 | 3564 | 2000 | Present | FI-like | CBDCA-PTX |
| FFPE-HGSOC-134 | High Grade Serous Ovarian Cancer | IIIC | 51,95 | 57 | 150 | 2012-02-09 | 2012-02-09 | 2014-02-21 | 2017-01-08 | 6028 | 350 | Present | FI-like | CBDCA |
| FFPE-HGSOC-135 | High Grade Serous Ovarian Cancer | IIIC | 46,44 | 53 | 163 | 2012-02-17 | 2012-02-17 | 2013-03-27 | 2016-09-26 | 3607 | 0 | Present | FI-like | CBDCA-PTX |
| FFPE-HGSOC-136 | High Grade Serous Ovarian Cancer | IIIC | 53,09 | 77 | 153 | 2012-02-27 | 2012-02-27 | 2012-11-29 | 2014-11-29 | 856 | 2000 | Absent | FI-like | CBDCA-PTX |
| FFPE-HGSOC-137 | High Grade Serous Ovarian Cancer | IIIC | 36,59 | 68 | 168 | 2012-03-09 | 2012-03-09 | 2013-05-30 | 2015-03-24 | 10579 | 6000 | Present | OSE-like | CBDCA-PTX |
| FFPE-HGSOC-138 | High Grade Serous Ovarian Cancer | IIIC | 67,77 | 78 | 163 | 2012-04-11 | 2012-04-11 | 2013-08-03 | 2018-02-20 | 81 | 0 | Absent | FI-like | CBDCA-PTX |
| FFPE-HGSOC-139 | High Grade Serous Ovarian Cancer | IIIC | 47,93 | 75 | 175 | 2012-04-27 | 2012-04-27 | 2014-03-04 | 2018-10-18 | 699 | 0 | Absent | OSE-like | CBDCA-PTX |
| FFPE-HGSOC-140 | High Grade Serous Ovarian Cancer | IV | 72,57 | 66 | 171 | 2012-05-09 | 2012-05-09 | 2013-01-18 | 2014-02-18 | 668 | 0 | Present | OSE-like | CBDCA-PTX |
| FFPE-HGSOC-141 | High Grade Serous Ovarian Cancer | IIIC | 70,94 | 70 | 160 | 2012-06-11 | 2012-06-11 | 2012-11-21 | 2018-04-13 | 1083 | 4800 | Present | OSE-like | CBDCA-PTX |
| FFPE-HGSOC-142 | High Grade Serous Ovarian Cancer | IIIC | 45,74 | 67 | 160 | 2012-06-18 | 2012-06-18 | 2013-11-11 | 2015-11-08 | 840 | 2200 | Present | OSE-like | CBDCA-PTX |
| FFPE-HGSOC-143 | High Grade Serous Ovarian Cancer | IIIC | 65,51 | 63 | 155 | 2012-07-10 | 2012-07-10 | 2013-01-07 | 2016-10-21 | 3687 | 2500 | Absent | OSE-like | Other |
| FFPE-HGSOC-144 | High Grade Serous Ovarian Cancer | IIIC | 56,76 | 61 | 156 | 2012-07-11 | 2012-07-11 | 2013-10-11 | 2017-05-11 | 954 | 3200 | Absent | OSE-like | CBDCA-PTX |
| FFPE-HGSOC-145 | High Grade Serous Ovarian Cancer | IV | 65,66 | 54 | 162 | 2012-07-24 | 2012-07-24 | 2013-06-22 | 2014-06-29 | 121 | 2000 | Absent | OSE-like | CBDCA-PTX |
| FFPE-HGSOC-146 | High Grade Serous Ovarian Cancer | IV | 68,84 | 46 | 157 | 2012-07-26 | 2012-07-26 | 2013-07-23 | 2018-01-10 | 15137 | 800 | Absent | OSE-like | CBDCA-PTX |
| FFPE-HGSOC-147 | High Grade Serous Ovarian Cancer | IIIC | 46,85 | 69 | 159 | 2012-08-30 | 2012-08-30 | 2013-07-15 | 2017-03-15 | 3728 | 7400 | Absent | FI-like | CBDCA-PTX |
| FFPE-HGSOC-148 | High Grade Serous Ovarian Cancer | IV | 66,98 | 82 | 165 | 2012-09-12 | 2012-09-12 | 2013-04-04 | 2017-09-20 | 595 | 500 | Absent | OSE-like | CBDCA-PTX |
| FFPE-HGSOC-149 | High Grade Serous Ovarian Cancer | IIIC | 63,48 | 65 | 160 | 2012-11-12 | 2012-11-12 | 2013-04-02 | 2016-10-19 | 3861 | 4800 | Absent | OSE-like | CBDCA-PTX |
| FFPE-HGSOC-150 | High Grade Serous Ovarian Cancer | IV | 42,96 | 64 | 165 | 2012-12-14 | 2012-12-14 | 2013-10-01 | 2014-07-25 | 344 | 2000 | Present | OSE-like | CBDCA-PTX |
| FFPE-HGSOC-151 | High Grade Serous Ovarian Cancer | IIIB | 41,54 | 63 | 167 | 2012-12-03 | 2012-12-21 | 2015-11-18 | 2018-10-19 | 35 | 20 | Absent | OSE-like | CBDCA-PTX |

**Supplementary Table 1. Description of the FFPE cohort's clinical data**
